# A shared alarmone-GTP switch underlies triggered and spontaneous persistence

**DOI:** 10.1101/2020.03.22.002139

**Authors:** Danny K. Fung, Jessica T. Barra, Jeremy W. Schroeder, David Ying, Jue D. Wang

**Affiliations:** Department of Bacteriology, University of Wisconsin-Madison, Madison, WI 53706, USA

**Keywords:** Antibiotic persistence, single cell reporter, phenotypic switch, metabolic switch, threshold, bet-hedging, ppGpp, GTP, spontaneous persistence, sublethal antibiotics

## Abstract

Phenotypically-switched, antibiotic-refractory persisters may prevent pathogen eradication. Although how triggered persistence via starvation-induced (p)ppGpp is well characterized, generation of persisters without starvation are poorly understood. Here we visualized the formation of spontaneous persisters in a small fraction of cells from growing wild type bacteria, revealing a striking single cell rapid switch from growth to dormancy. This switch-like entrance is triggered by GTP dropping beneath a threshold due to stochastic production and self-amplification of (p)ppGpp via allosteric enzyme activation. In addition, persisters are induced by lethal and sublethal concentrations of cell wall antibiotics by inducing (p)ppGpp via cell wall stress response. Thus spontaneous, triggered and antibiotic-induced persisters can all stem from a common metabolic switch: GTP depletion by (p)ppGpp induction, and each pathway of persister formation is activated by different (p)ppGpp synthetases. These persistence pathways are likely conserved in pathogens which may be exploited to potentiate antibiotic efficacy.

## Introduction

Persisters are phenotypically switched, multidrug-refractory cells in a genetically susceptible population that are able to survive lethal doses of antibiotics, while preserving the ability to resume growth when treatment is discontinued (Balaban et al., 2004; Bigger, 1944; Harms et al., 2016). Persisters do not possess heritable antibiotic resistance traits, yet their survival may contribute to the evolution of antibiotic resistance, which ultimately leads to antimicrobial failure (Levin-Reisman et al., 2017).

Persister cells exist in natural populations at low frequency, making their identification challenging and understanding the mechanisms of their formation even more difficult. Persisters are classified as triggered (Type I) persistence, which can be triggered by starvation and are enriched in stationary phase or starved populations; and spontaneous (Type II) persisters, which are generated as a rare fraction during balanced growth (Balaban et al., 2004). Triggered persistence has been proposed to be generated via the starvation-inducible nucleotide (p)ppGpp (Korch et al., 2003; Nguyen et al., 2011). Alternatively, it is argued that triggered persisters do not form due to (p)ppGpp but via depletion of ATP (Conlon et al., 2016). Even less is known about spontaneous persisters because direct observation of stochastic entrance into persistence in wild-type cells has not been reported. Although overexpressing (Kaspy et al., 2013; Rotem et al., 2010) or mutating (Balaban et al., 2004; Moyed and Bertrand, 1983) toxin-antitoxin (TA) modules can increase spontaneous persister formation, the physiological role of TA modules in persistence of wild-type bacteria has been recently challenged (Harms et al., 2017; Shan et al., 2017).

Apart from arising prior to antibiotic assault, persisters can also form in response to antibiotic treatment. However, it is difficult to differentiate drug-induced persistence from spontaneous persistence, drug-induced tolerance, specific transcriptional response to each class of drugs or byproduct of cellular damage from drug treatment (Balaban et al., 2019; Johnson and Levin, 2013). The mechanism of genesis and metabolic traits of antibiotic-induced persistence remain poorly understood.

Here we characterized the pathways of persister formation in the Gram-positive bacterium *Bacillus subtilis* and showed that triggered and spontaneous persisters, as well as antibiotic-induced persisters, can all stem from a common metabolic switch in Gram-positive bacteria: GTP depletion by (p)ppGpp induction. We report the first direct recording of spontaneous persister entrance dynamics in single cells, revealing a striking switch from active growth to dormancy, and reveal (p)ppGpp as the key to the switch-like dynamics. Finally, we pinpoint the different (p)ppGpp synthetases that are responsible for different persister pathways. This core mechanism of persister formation that we elucidated may be conserved in other bacteria including pathogens.

## Results

### (p)ppGpp mediates both triggered and spontaneous persistence

Antibiotic persistence is the phenomenon where a fraction of cells in a population survive intensive and prolonged bactericidal antibiotic treatment: at concentrations many times above the minimal growth-inhibitory concentrations (MIC) (Balaban et al., 2019). To characterize antibiotic persistence of the Gram-positive bacterium *B. subtilis*, we first determined the MICs of *B. subtilis* wild-type strain NCIB 3610 (Fig 1A and S1A), then treated exponentially growing cells with cell wall-damaging antibiotic vancomycin (VAN) at a range of concentrations multiple times (4-fold to 40-fold) above its MIC (Fig 1B). We observed rapid killing of the bulk population, followed by an extended period of survival of a small fraction of cells (∼0.1%). This bimodal killing and prolonged surviving population follow the definition of persisters. The size of the surviving fraction was independent of drug concentrations, and the surviving cells were sensitive to new treatment of antibiotics upon regrowth (Fig 1C), confirming that they are due to phenotypically switched persistence rather than genetically altered resistance, heteroresistance, or tolerance (Balaban et al., 2019; Bigger, 1944; El-Halfawy and Valvano, 2015). Similar results were obtained from treatment by DNA damaging antibiotic ciprofloxacin (CIP) (Fig S1B and S1C) except for decreased survival of cells in ciprofloxacin (∼0.01%) compared to vancomycin (∼0.1%). We found this decrease is due to increased cell lysis triggered by resident prophages upon ciprofloxacin treatment (Goranov et al., 2006). When we applied ciprofloxacin to a strain deleted of prophages PBSX and SPβ, endpoint survival increased to ∼0.1% (Fig 1D) similar to that of VAN, suggesting that the persister population can tolerate different antibiotics.

**Figure 1.**
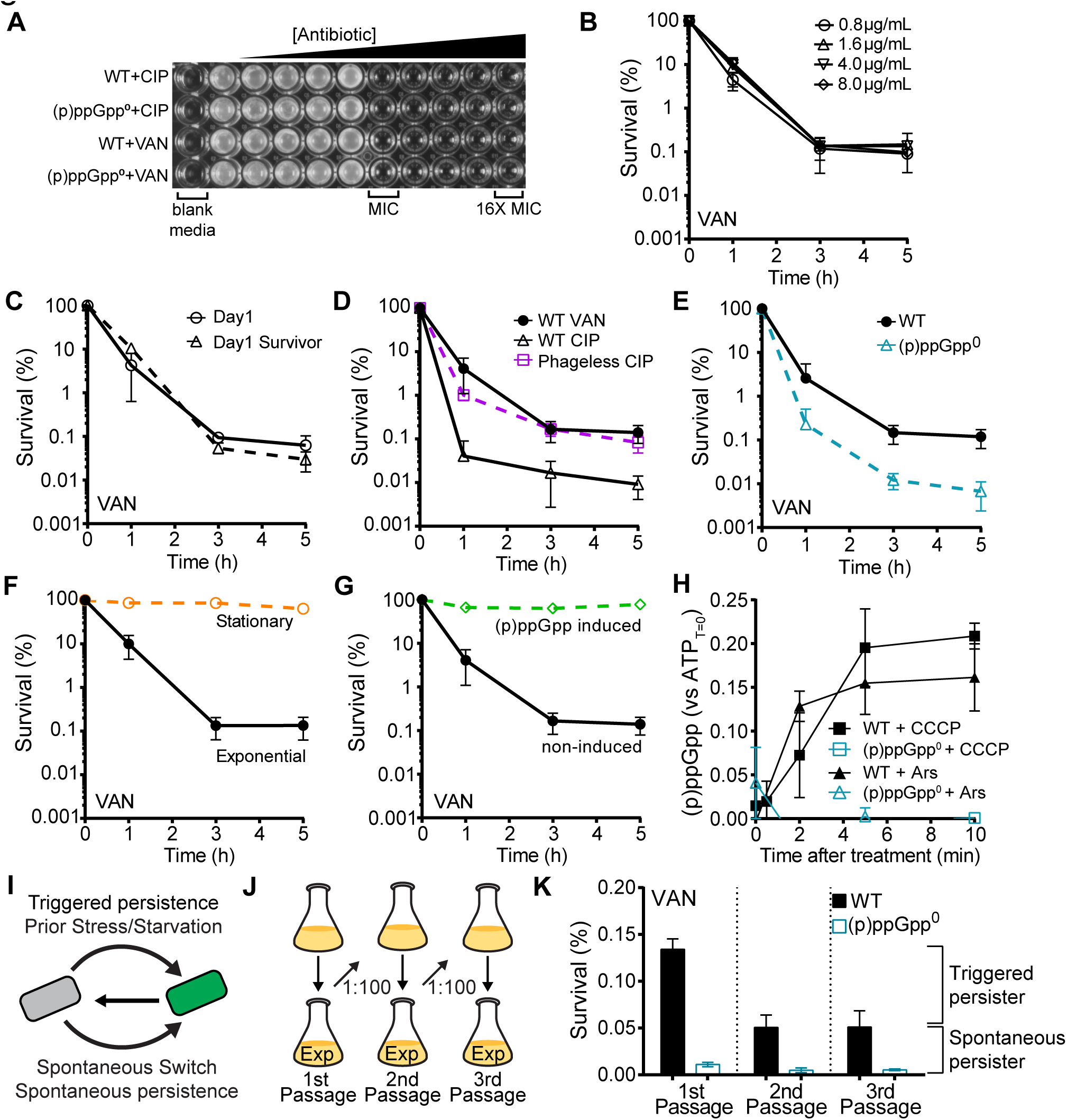
(p)ppGpp mediates persistence to antibiotics. (A) (p)ppGpp does not affect antibiotic sensitivity. Minimal inhibitory concentrations (MICs) of WT and (p)ppGpp^0^ to VAN and CIP. See also Fig S1A. (B) Persistence is independent of antibiotic concentration. Survival of WT cells to concentrations of VAN ranging from 4x to 40x MIC for up to 5 h. See also Fig S1B. (C) Persistence is not due to rare mutants. Survival of exponentially-growing WT cells (Day 1, solid lines), or its successor population derived from survivors after 5 h treatment (Day 1 survivor, dashed lines) to VAN for up to 5h. See also Fig S1C. (D) Persisters are multidrug tolerant. Survival of WT and prophage-cured mutant (Δ*zpdN* ΔSPβ Δ PBSX) to VAN or CIP treatment. (E) Persistence is dependent on (p)ppGpp. Survival of WT (solid lines) and (p)ppGpp^0^ mutant (dashed lines) to VAN for 5 h. See also Fig S1D and E. (F to H) Stationary phase or (p)ppGpp accumulation induced by starvation induce majority of cells into persisters. (F) Survival of exponentially growing or stationary phase WT cells to VAN. (G) Survival of (p)ppGpp-induced or non-induced WT cells to VAN. (p)ppGpp induction was done by treatment with amino acid analog for 30 min. See also Fig S1F. All data points represents mean ± SD, n ≥ 3. (H) (p)p-pGpp is induced by inhibition of ATP synthesis. WT and (p)ppGpp^0^ cells were treated with CCCP or arsenate and measured for changes in (p)ppGpp with TLC over time as indicated. Values represent mean ± SD, n = 2. (I) Persistence can be due to the presence of triggered (Type I) persisters or spontaneous (Type II) persisters. (J and K) (p)ppGpp mediates both triggered and spontaneous persistence. (J) WT and (p)ppGpp^0^ cells were grown to log phase (1st passage) and sub-cultured by 100-fold for two additional rounds (2nd passage and 3rd passage). (K) Different passages were tested for survival to VAN for 5 h. Persisters which were eliminated by serial passage were triggered persisters, while spontaneous persisters remained. Values represent mean ± SD, n = 5.

We found that persister formation depends on the nucleotide (p)ppGpp. Despite having similar MICs to antibiotics (Fig 1A and S1A), a mutant devoid of all (p)ppGpp synthetases that cannot produce (p)ppGpp [(p)ppGpp^0^] has a significantly (∼ 10-fold) reduced persister fraction (Fig 1E, S1D and S1E). Furthermore, conditions that induce (p)ppGpp, including stationary phase, amino acid starvation (Kriel et al., 2012), or growth transition from rich to amino acids-depleted media all resulted in dramatic enrichment of persisters: 30%-80% of the total population of cells can survive antibiotics for prolonged hours (Fig 1F, 1G and S1F). Lastly, we found that chemical inhibition of ATP synthesis, which promotes persistence in *B. subtilis* (Fig S1G) as previously shown for *S. aureus* (Conlon et al., 2016), can also induce (p)ppGpp (Fig 1H).

Persisters have been categorized as two types based on how they are generated: triggered persisters, which are triggered by starvation and upon entrance to stationary phase, or spontaneous persisters (Fig 1I). To determine the category of persisters in growing cell populations, we performed passage experiments in which cells were subjected to repeated sub-culturing in early exponential phase (OD_600_ < 0.2) to diminish triggered persisters (Fig 1J). The fraction of persisters dropped from ∼0.1% to ∼0.05% after the first passage but remained steady after the second passage. This indicates there are ∼0.05% triggered persisters in the first generation and ∼0.05% spontaneous persisters continuously generated during balanced growth in successive passages (Fig 1K, black bars). Importantly, the (p)ppGpp^0^ mutant exhibited more than ten-fold reduction of persisters relative to wild-type in both the first generation and successive passages (Fig 1K, blue bars), indicating that (p)ppGpp is a major source of both starvation-triggered persisters and spontaneous persisters.

### GTP depletion is the metabolic effector of (p)ppGpp-mediated persistence

(p)ppGpp accumulation has multiple downstream effects in bacteria (Haugen et al., 2008; Liu et al., 2015; Potrykus and Cashel, 2008). These include transcription regulation which is largely dependent on the transcription factor CodY in *B. subtilis* (Kriel et al., 2014; Sonenshein, 2007) and developmental processes including competence (Hahn et al., 2015) or sporulation (Ochi et al., 1982) in *B. subtilis*. In addition, Type II toxin-antitoxin systems were implicated in persister formation in a (p)ppGpp dependent manner (Kaspy et al., 2013). However, we found that deletion of *codY* (Fig 2A), the three Type II Toxin-Antitoxin systems (Δ3TA) (Fig 2B), or the competence regulator *comK* (Fig 2C) had no effect on persistence. Disruption of the sporulation pathway (Δ (p)ppGpp-independent and drug-specific effect on starvation-triggered persistence (Fig 2D), but no effect on spontaneous persistence (Fig 2E and S2A). These altogether suggest that persistence induced by (p)ppGpp is unlikely to be a result of transcriptional changes, activity of TA systems, or specific (p)ppGpp-related developmental processes in *B. subtilis*.

**Figure 2.**
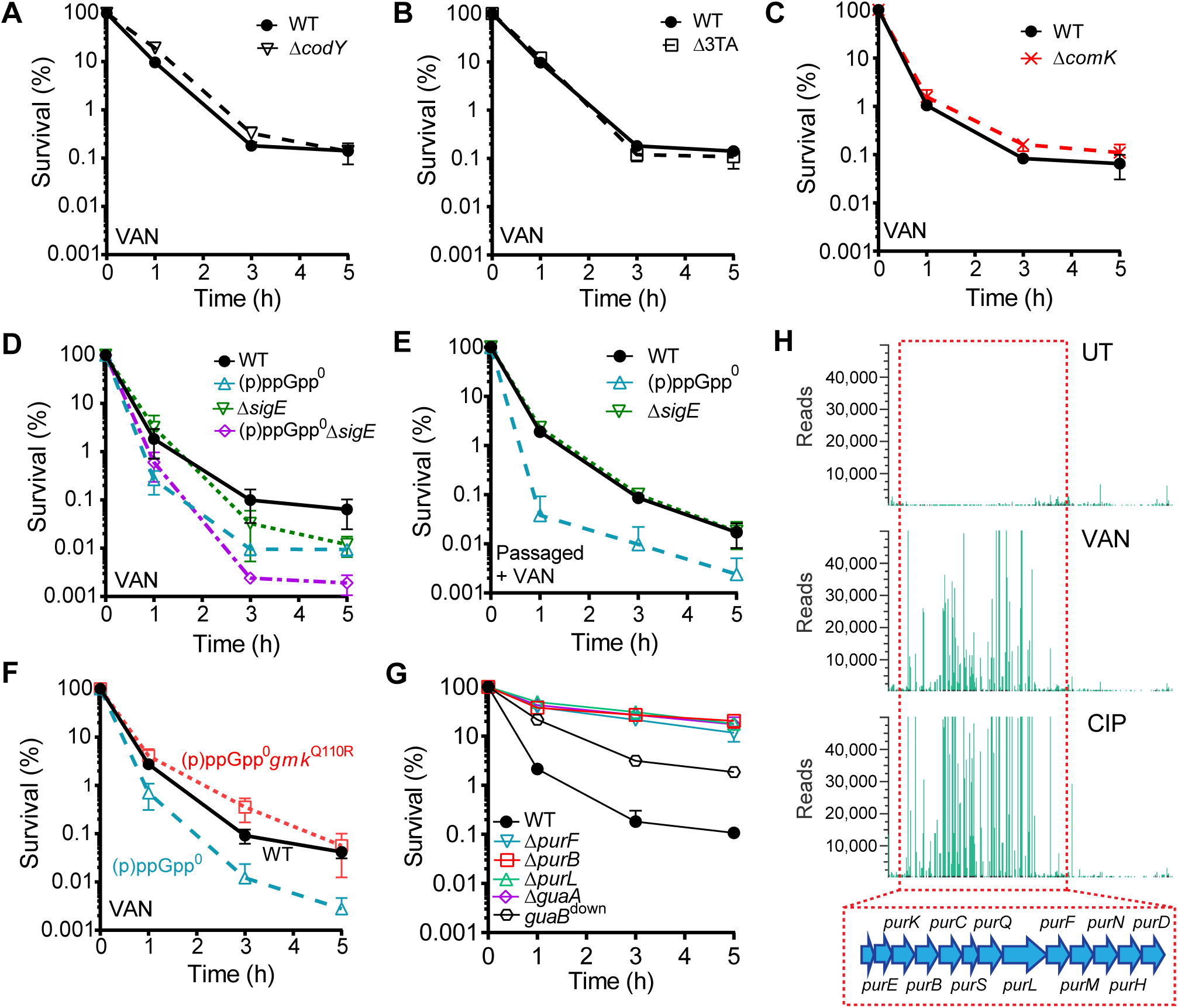
Persistence by (p)ppGpp-mediated GTP down-regulation. (A to E) Persistence through (p)p-pGpp is not due to transcription regulation, TA systems, competence or sporulation. Survival of exponentially-growing (A) WT and Δ*codY* cells, (B) WT and a mutant lacking all known TA systems (Δ3TA), (C) WT and Δ*comK* cells, and (D) WT, (p)ppGpp^0^, Δ*sigE,* and (p)ppGpp^0^ Δ*sigE* cells to VAN treatment for 5h. (E) WT, (p)ppGpp^0^ and Δ*sigE* cells in passaged population to VAN treatment for 5h. Values represent mean ± SD, n = 3. See also Fig S2A. (F) Lowering GTP in (p)ppGpp^0^ restores persistence. Survival of WT (black), (p)ppGpp^0^ (blue), and (p)ppGpp^0^ *gmk*^Q110R^ (red) to VAN treatment for 5 h. Values represent mean ± SD, n = 3. See also Fig S2B. (G) Depleting GTP in (p)ppGpp+ cells increases persistence. Survival of WT, Δ*purB*, Δ*purL*, Δ*purF*, Δ*guaA*, and GuaB depleted strain (*guaB*^Down^) to VAN treatment. Values represent mean ± SD, n = 3. (H) Enrichment of *de novo* purine biosynthesis mutants in genome-wide screening for tolerance mutants. Location and frequency of transposon insertions in the *pur* operon (red box) before (UT) and after VAN or CIP treatment from Tn-Seq experiment. See also Fig S2D.

We previously showed that (p)ppGpp directly inhibits enzymes (e.g., the GDP synthetase Gmk) in GTP synthesis pathways (Fig S2C) and robustly depletes GTP levels in *B. subtilis* (Kriel et al., 2012; Liu et al., 2015). Correspondingly, the (p)ppGpp^0^ strain has elevated GTP levels. To test whether (p)ppGpp enables persister formation by reducing GTP levels, we obtained a mutant (*gmk*^Q110R^, see methods) that reduces GTP levels of the (p)ppGpp^0^ strain back to wild-type levels (Fig S2B). We found that the (p)ppGpp^0^ *gmk*^Q110R^ mutant had restored persister level back to wild-type (Fig 2F), indicating that GTP depletion is a persistence effector downstream of (p)ppGpp.

Furthermore, in (p)ppGpp^+^ cells, lowering GTP synthesis by depleting the transcription of *guaB* (encoding IMP dehydrogenase that generates the GTP precursor XMP) further increased persistence by ∼10-fold from wild-type levels (∼0.1% to ∼1%) (Fig 2G), indicating GTP depletion is sufficient to generate higher fractions of persisters even in the absence of starvation.

Finally, we performed genome-wide transposon sequencing (Tn-Seq) analysis in wild-type *B. subtilis* to identify genes whose disruption leads to increased persistence. The strongest hits mapped to more than ten genes involved in purine biosynthesis. In particular, insertions in the *pur* operon and the gene *guaA* (GMP synthase) displayed more than 500-fold enrichment in relative abundance after antibiotic treatment (Fig 2H and S2D) and increased persistence (Fig 2G), indicating that disrupting synthesis of precursors of GTP aids antibiotic survival. These data altogether demonstrate that GTP depletion is a key effector of persister formation.

### Single-cell GTP depletion dictates persistence

Next, we examined whether persisters stem from overall low GTP in all cells, or stem from rare low-GTP cells due to heterogeneity of GTP levels between individual cells. To visualize GTP levels in single cells, we took advantage of our previous finding that depletion of GTP results in strong activation of the *ilvB* promoter (P*_ilvB_*) (Handke et al., 2008; Kriel et al., 2014), and constructed a single-cell fluorescent reporter transcriptionally fused to P*_ilvB_.* The reporter sensitively exhibited dose-dependent responses to differences in cellular GTP levels (Fig 3A). Its expression was also highly responsive to conditions that lead to GTP depletion including (p)ppGpp accumulation at stationary phase (Fig 3B), carbon or amino acid starvation (Fig 3C and S3A), or treatment with ATP synthesis inhibitors (Fig 1H and S3B). The P*_ilvB_* activity is also reciprocal to the activity of the ribosomal P1 promoter (Fig S3C) which is reported to be inhibited by reduced GTP (Krásný and Gourse, 2004).

**Figure 3.**
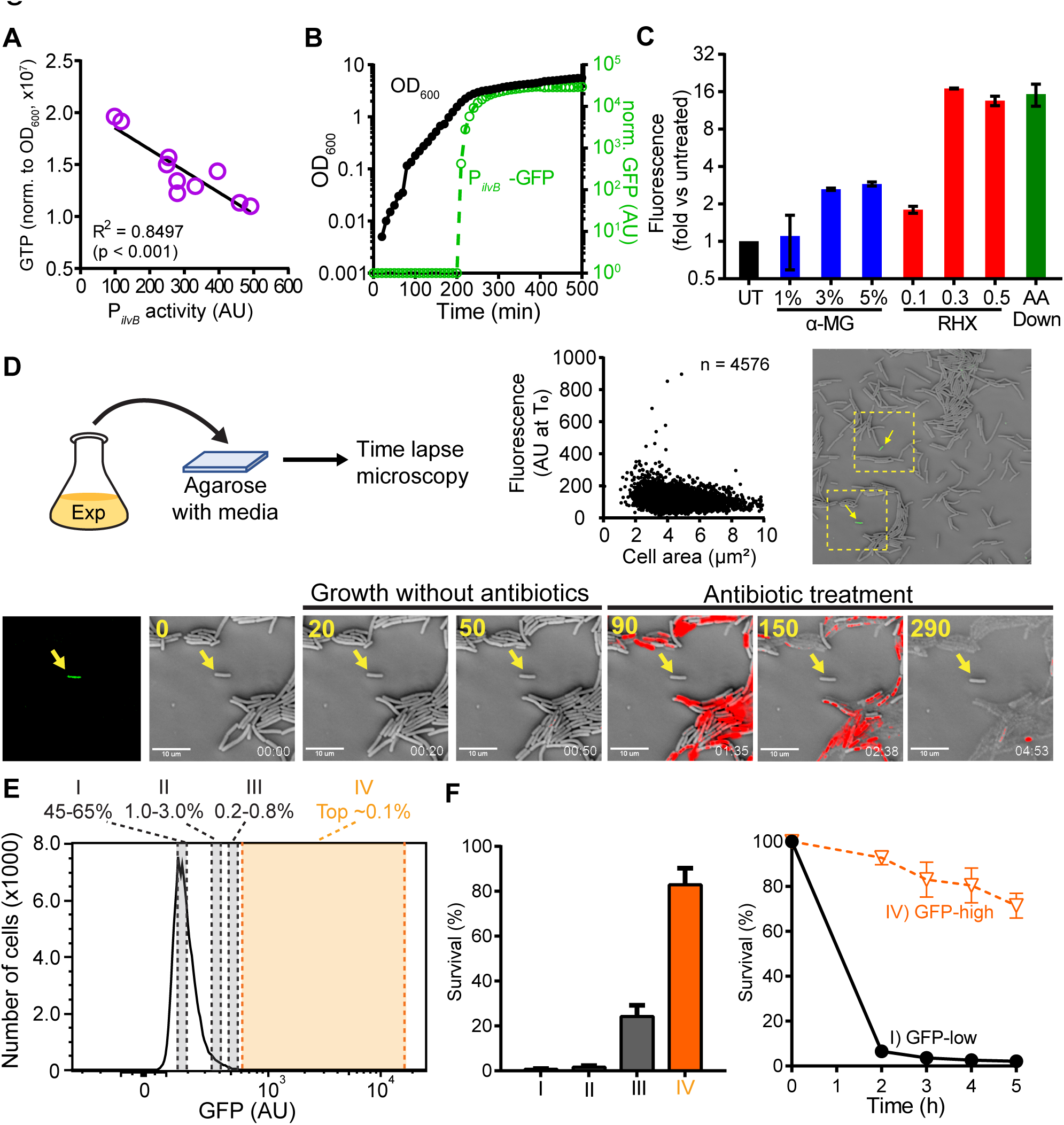
Single-cell GTP depletion underlies persistence. (A to C) Application of single-cell fluorescent reporter to monitor GTP depletion. (A) P*_ilvB_* activity inversely correlates with cellular GTP levels. GTP levels in a panel of *guaB* mutants (Bittner et al 2014) were measured by TLC and P*_ilvB_* activity were measured by fluorescence microscopy. n = 3 for TLC and n > 200 cells for microscopy. Solid line indicates linear regression. (B) Changes in P*_ilvB_*-*gfp* (green) and OD_600_ (black) during growth into stationary phase. GFP signals were normalized to OD_600_. (C) WT cells were treated with variable concentrations of alpha-methylglucoside (α-MG, blue), arginine hydroxamate (RHX, red), or amino acids down-shift (AA down, green). Single-cell fluorescence was measured after 2 h using fluorescence microscopy. Values represent mean ± SD from n > 200 cells of two independent experiments for each condition. UT represents untreated control. (D to F) Identification and antibiotic tolerance of low-GTP persisters. (D) Cells containing the P*_ilvB_* reporter were grown in liquid media to exponential phase, then patched on agarose pads made with growth media for analysis of GTP levels in single cells with microscopy. GTP-depleted cells were identified (arrow) and monitored for antibiotic survival using carbenicillin and Sytox-blue staining (red) for viability. Values in yellow indicate time in min. (E) FACS sorting of exponentially-growing WT cells into fractions with decreasing GTP levels (I to IV) based on P*_ilvB_* activities as indicated. (F) Survival to VAN of sorted populations from (E). Values represent mean ± SD, n = 3.

Using the P*_ilvB_* reporter, we examined GTP levels in individual cells from exponentially-growing cultures with time-lapse microscopy (Fig 3D). We found that while the vast majority of cells displayed basal fluorescence indicative of high GTP, a few cells (< 0.1%) displayed bright fluorescence indicative of low GTP levels (Fig 3D). Cells that had high P*_ilvB_* activities (low GTP) were non-growing, while the rest of the cells maintained growth and division (Fig 3D, 0 min to 50 min). We subsequently treated cells with the antibiotic carbenicillin and tracked cell survival (Fig 3D, 90 min to 290 min, Movie S1). As expected, most cells were rapidly killed as indicated by permeability to Sytox blue (red) and eventually lysed. However, the low-GTP cells remained viable as indicated by lack of lysis and Sytox blue staining (Fig 3D arrow, 90 min to 290 min), suggesting that they are refractory to antibiotic treatment.

To test whether these low GTP cells were *bona fide* persisters that can regrow after removal of antibiotics, we used fluorescence-activated cell sorting (FACS) to isolate cells with low GTP (high P*_ilvB_* activity) from the bulk population. Using flow cytometry, we observed a small fraction (∼0.1%) of high P*_ilvB_* activity cells (∼3-fold above that of the bulk population) in exponentially growing cultures that constitute a heavy-tailed distribution (Fig 3E). We sorted the populations according to their fluorescence using FACS, then separately treated with vancomycin. Strikingly, ∼80% of high P*_ilvB_* activity cells (the top ∼0.1%) were able to survive antibiotic treatment and form colonies upon drug-removal (Fig 3F), suggesting that nearly all purified cells with low levels of GTP are antibiotic-refractory persisters. In addition, the size of this subpopulation increases in response to amino acid starvation with a proportional increase in persistence (Fig S3D to F), suggesting that the reporter allows detection of persister dynamics in the population.

### Time-lapse observation of spontaneous persister entrance reveals that (p)ppGpp enables switch-like dynamics from growth to persistence

Our P*_ilvB_* reporter enables us to monitor the dynamics of the spontaneous entrance to persistence. We seeded highly diluted, exponentially growing cells containing the reporter on agarose pads made with growth media (Fig 4A) and monitored their growth with time-lapse microscopy (Fig 4B). We followed ∼30,000 growth and division events, which allowed us to identify 10 cells spontaneously entering persistence from active growth (Fig 4C, Movies S2) before the microcolony reached high cell density (Fig S4A). Thus, spontaneous persister entrance is observed in wild-type bacterial cells. Importantly, we can now retrospectively examine their trajectory before entering persistence by analyzing the dynamics of their growth, division, and changes in P*_ilvB_* activity over time (Fig 4E and S4B).

**Figure 4.**
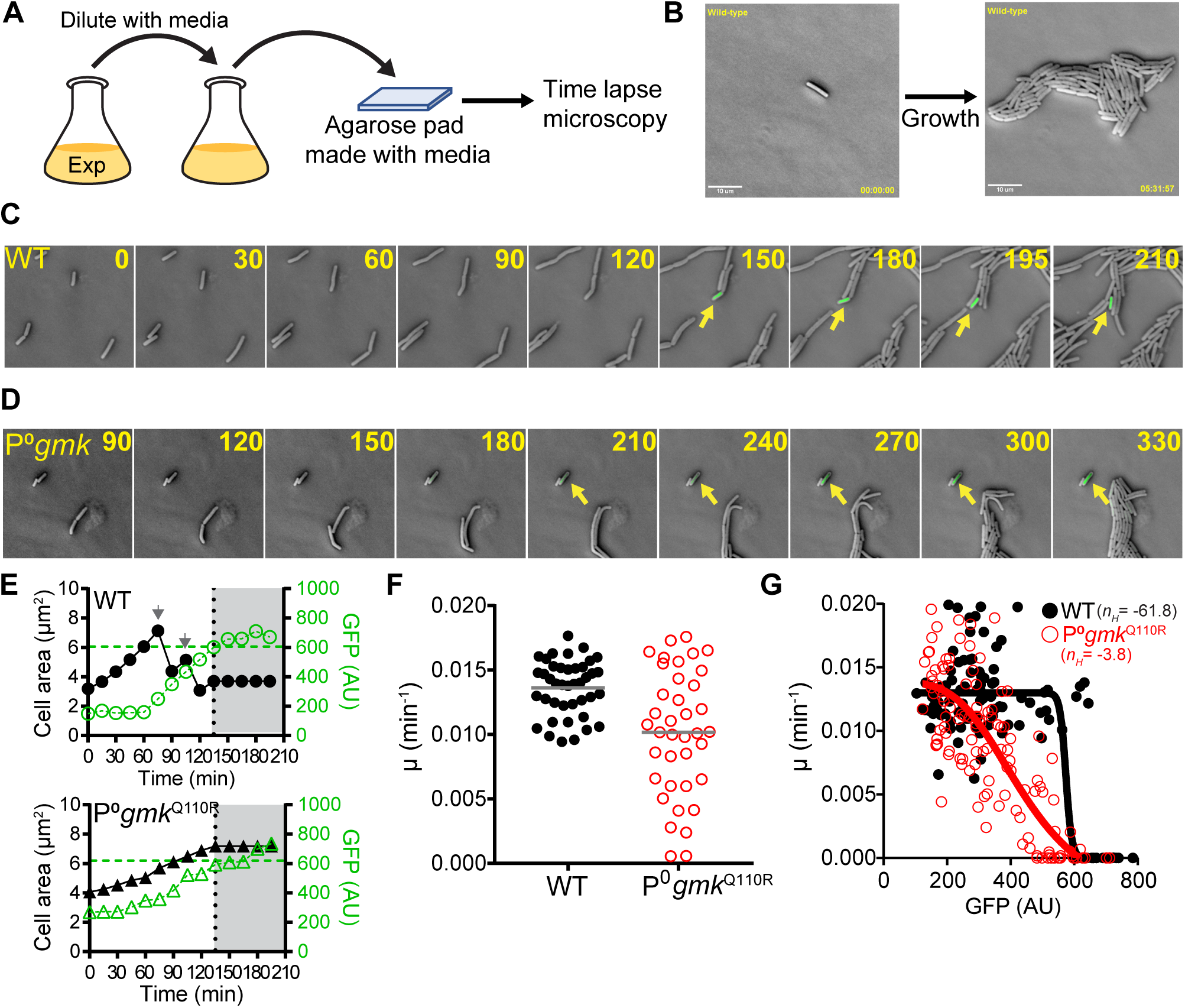
GTP depletion by (p)ppGpp leads to rapid persister formation. (A) Schematic of timelapse microscopy to monitor growth and persister formation. Cells containing the P*_ilvB_* reporter were grown in liquid media to exponential phase, then diluted and patched on agarose pads made with growth media. Microcolony growth is monitored with timelapse microscopy. An example is provided in (B). (C and D) Formation of spontaneous persisters during active cell growth in (C) WT and (D) (p)ppGpp^0^ *gmk*^Q110R^ mutant. Representative time-lapse images of persister formation are shown. Numbers indicate time in min. See also Fig S4A. (E) Changes in P*_ilvB_* activity (green) and cell size (black) during stochastic entrance into persistence in WT and (p)ppGpp^0^ *gmk*^Q110R^ mutant. See also Fig S4B and C. Persistence entry was marked by vertical dotted lines and horizontal dashed lines. Arrows indicate cell division. (F) Variations in specific growth rates (µ) of growing wild-type (WT, solid black circles) and (p)ppGpp^0^ *gmk*^Q110R^ cells (P^0^*gmk*^Q110R^, open red circles). Grey bars indicate the mean. n > 40 cell tracks for each. (G) Correlation of specific growth rate (µ) and P*ilvB* activities in WT (solid black circles) and (p)ppGpp^0^ *gmk*^Q110R^ mutant (open red circles) during persistence entrance. Solid (WT) and dashed lines ((p)p-pGpp^0^ *gmk*^Q110R^) indicate non-linear regression (variable slope) model fitted to either population. *n*H represents the Hill slope.

Through this analysis, we observe a striking switch-like single cell growth pattern in persister transition. Despite a gradual increase of P*_ilvB_* activity before entering persistence, each pre-persister cell maintained the same elongation rate as actively growing cells. This is followed by a sudden cessation of growth once the P*_ilvB_* activity reached a threshold (Fig 4E and S4B). The P*_ilvB_* activity thresholds were highly similar between persister entrance events in different cells (Fig 4E, 4G and S4B), suggesting that switch-like entrance to persistence is triggered by a similar GTP level.

To examine the respective roles of GTP and (p)ppGpp on persister dynamics, we also examined persister entrance of the (p)ppGpp^0^ *gmk*^Q110R^ mutant, which has similar GTP levels and persister fractions as wild-type cells (Fig 2D) but without (p)ppGpp. We found that a few cells also formed spontaneous persisters at a similar reporter threshold, indicating that low GTP dictates persister formation regardless of (p)ppGpp (Fig 4D, 4E and S4C). However, there is a drastic difference in the persister entrance dynamics in the (p)ppGpp^0^ *gmk*^Q110R^ mutant compared to wild-type. The switch-like dynamics exhibited in wild-type cells were abolished in the mutant, with cells first slowing down growth as P*_ilvB_* activity increases and gradually entering dormancy (Fig 4D and 4E). The GTP levels and growth rate in the (p)ppGpp^0^ *gmk*^Q110R^ mutant exhibit a linear correlation, in strong contrast to a switch-like relationship in wild-type (Fig 4F and 4G, black vs red). Therefore, (p)ppGpp not only indirectly enables persistence through GTP depletion but is also responsible for the rapid switch dynamics in spontaneous persistence.

### Cell-wall antibiotic treatments promote persister formation through (p)ppGpp

Apart from starvation-triggered and spontaneous persisters which both arise prior to antibiotic treatment, we found a strong increase of persister formation triggered by antibiotic treatment. We found that P*_ilvB_* activity increases in all cells after they are exposed to cell wall antibiotics, including carbenicillin, vancomycin, and bacitracin (Fig 5A and 5C). In addition, although most cells were killed by the antibiotics, a minor fraction of cells (∼10%) became dormant and were not lysed during antibiotic treatment (Fig 5B, Carbenicillin treatment). Most of the dormant cells were unable to regrow when the antibiotics was removed, presumably due to unrecoverable cellular damage. However, a few of them were able to resume growth, suggesting that they were persisters (Fig 5B, Antibiotic removal). This observation raises the intriguing possibility that antibiotic treatment itself can induce persisters via the same GTP pathway.

**Figure 5.**
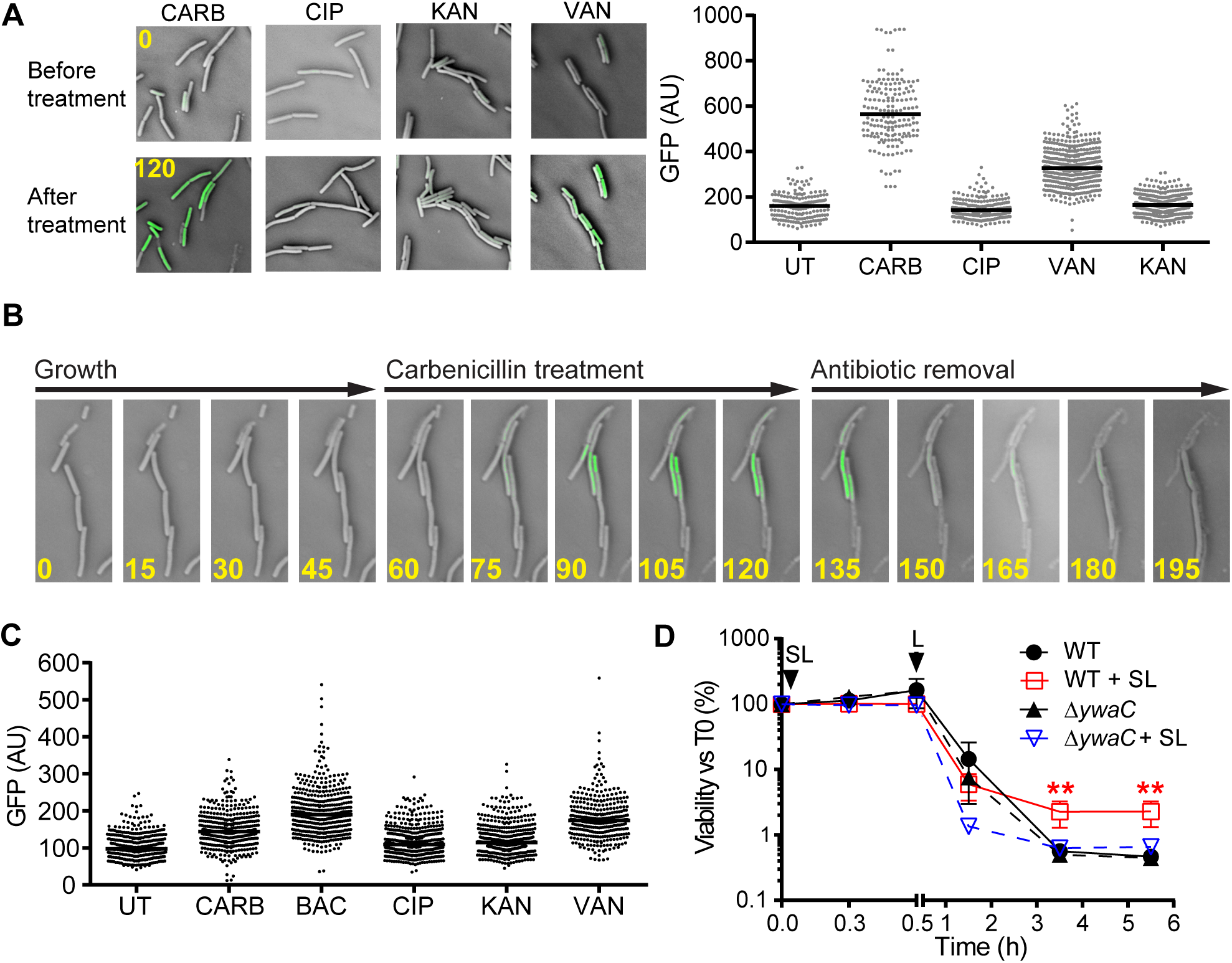
(p)ppGpp accumulation in response to specific antibiotics enables induced persistence. (A) Cell wall antibiotics induce (p)ppGpp accumulation. WT cells containing P*_ilvB_* reporter were grown without antibiotic followed by treatment with lethal concentrations of carbenicillin (CARB), ciprofloxacin (CIP), kanamycin (KAN), or vancomycin (VAN). Numbers indicate time in minutes. (B) Time-lapse images of antibiotic-induced survival upon antibiotic treatment. WT cells containing the P*_ilvB_* reporter (green) were allowed to grow in the absence of antibiotic followed by carbenicillin treatment. Antibiotic was removed by penicillinase treatment. Numbers indicate time in min. (C) WT cells containing P*_ilvB_* reporter were exposed to sublethal (0.5x MIC) bacitracin (BAC) and various antibiotics as indicated for 30 min. UT: no treatment. (D) Sub-lethal cell wall antibiotic increases persistence. WT or Δ*ywaC* cells were subjected to sublethal (SL) BAC or without treatment for 30 min, followed by exposure to lethal BAC treatment (L) for up to 5 h. Values represent mean ± SD, n = 3. **p < 0.005 (Student’s t test).

Interestingly, we found that GTP-mediated persisters can be triggered by not only lethal concentrations (> MIC) but also sublethal concentrations (< MIC) of antibiotics. We treated cells with 0.5x MIC concentration of bacitracin and found that although this treatment does not trigger cell death, it leads to significant induction of P*_ilvB_* activity (Fig 5C). Importantly, when followed by lethal bacitracin treatment (3x MIC) (Fig 5D), we found that sublethal pretreated cells have much higher numbers (∼5-fold) of survivors (Fig 5D) compared to non-pretreated cells. Therefore, generation of persisters due to sublethal antibiotic exposure may be an adaptive response that strongly promotes cell survival against subsequent lethal antibiotic assault.

### Different (p)ppGpp synthetases are responsible for different pathways of persister formation

(p)ppGpp can be synthesized by three synthetases in *B. subtilis*: the bifunctional enzyme RelA and the small alarmone synthetases YjbM (SAS1) and YwaC (SAS2) (Nanamiya et al., 2008; Srivatsan et al., 2008). First, we found that the formation of triggered persisters due to amino acid starvation (Fig 1G and S1F) is mediated by RelA, since a RelA synthetase-defective mutant (*relA^Syn^*) has abolished starvation-triggered persistence (Fig 6A). In addition, we found that RelA is also responsible for (p)ppGpp synthesis upon treatment with ATP synthesis inhibitor (Fig S5A and S5B), a condition that promotes triggered persister formation (Conlon et al., 2016). Thus, starvation-triggered persisters are mainly generated by RelA-dependent (p)ppGpp synthesis.

**Figure 6.**
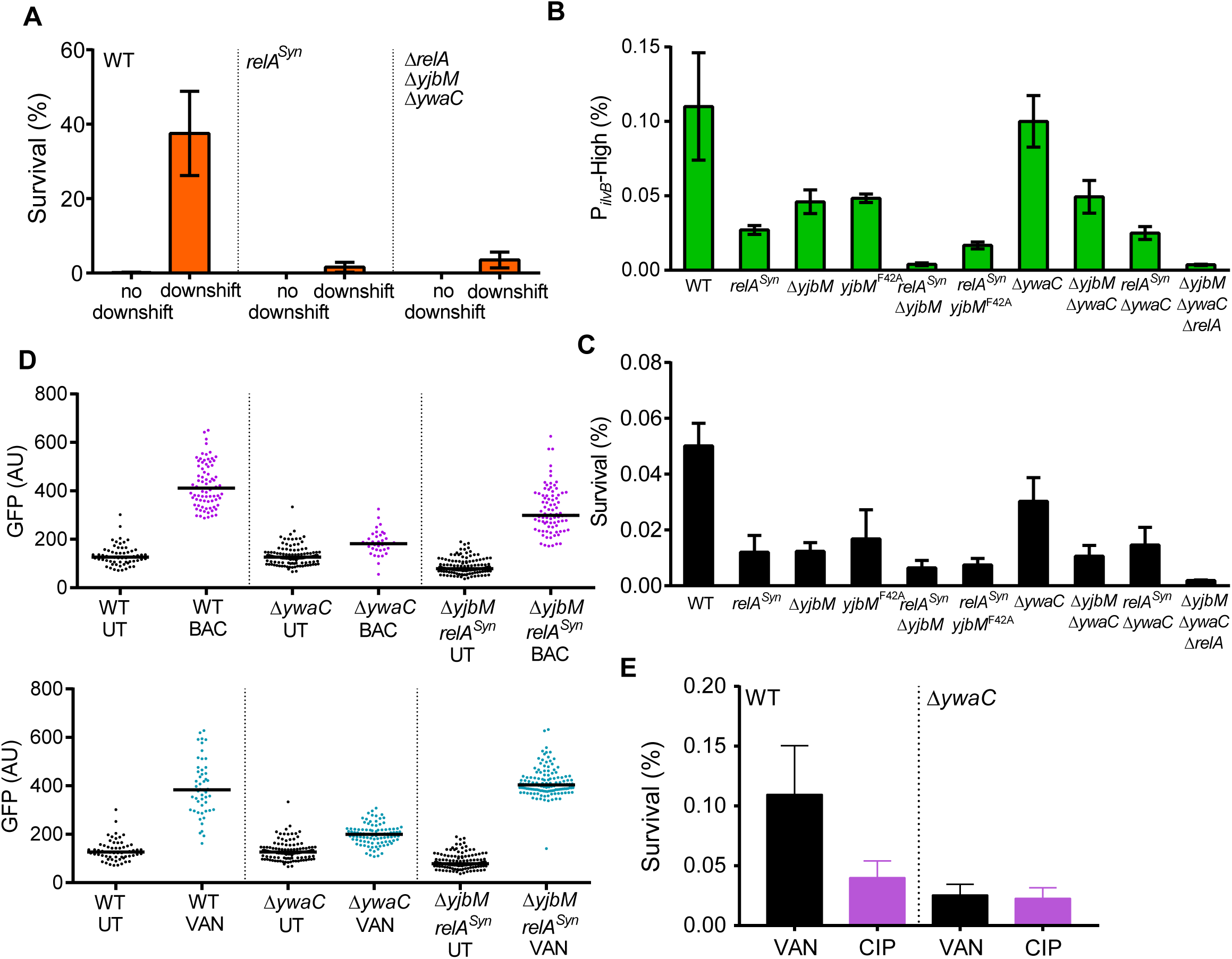
Different (p)ppGpp synthetases mediate different pathways to persistence. (A) (p)ppGpp synthesis by RelA mediates triggered persistence. WT, *relA^Syn^* (synthetase inactive), and (p)ppGpp^0^ cells were grown to exponential phase followed by nutrient downshift from rich media to amino acids depleted media for 30 min. Cells were then subjected to VAN treatment for 3 h. Values represent mean ± SD, n = 3. (B and C) Synergistic (p)ppGpp synthesis by RelA and YjbM leads to spontaneous persistence. (B) Frequency of cells with high P*_ilvB_* activity in WT and (p)ppGpp biosynthesis mutant populations (see also Fig S5C and S6). Cells were grown to and passaged at exponential phase followed by analysis by flow cytometry (n > 10^6^ cells each, three independent replicates). (C) Survival of populations from (B) to VAN treatment for 5 h. Values represent mean ± SD, n = 3. (D and E) Cell wall antibiotic induces (p)ppGpp through YwaC to enable adaptive persistence. (D) Cell wall antibiotic induces (p)ppGpp through YwaC. WT, Δ*ywaC* and Δ*yjbM relA^Syn^* cells containing the P*ilvB* reporter were allowed to grow in the absence of antibiotic, followed by BAC or VAN treatment for 1h. UT: Before treatment. (E) YwaC promotes survival to cell wall antibiotic. WT and Δ*ywaC* populations were subjected to VAN or CIP treatment for 5 h. Values represent mean ± SD, n = 3.

Next, we identified two (p)ppGpp synthetases responsible for spontaneous persistence: RelA and the small synthetase YjbM. To study exclusively spontaneous persisters, we analyzed passaged cell populations devoid of triggered persisters (Fig S5C and S5D, Fig S6). We found that both *relA^Syn^* or Δ*yjbM* mutants showed significant reductions (∼5-fold or ∼2-fold respectively) of the spontaneous persister population (Fig 6B), whereas *relA^Syn^* Δ*yjbM* double mutant almost completely eliminated spontaneous persisters similar to the (p)ppGpp^0^ mutant (Fig 6B, 6C and S5E), indicating a synergistic role of *relA* and *yjbM* on the formation of spontaneous persisters.

YjbM forms a homotetramer and its ability to synthesize ppGpp is allosterically activated by pppGpp (Steinchen et al., 2015). We constructed a *yjbM* mutant containing an alanine substitution (YjbM^F42A^) that disrupts the allosteric pppGpp binding site without affecting its active site. The *yjbM*^F42A^ mutant showed reduced spontaneous Δ*yjbM* mutant (Fig 6B and 6C), suggesting that allosteric activation of YjbM by pppGpp allows the formation of spontaneous persisters through amplification in (p)ppGpp synthesis.

Lastly, we found that (p)ppGpp accumulation in response to cell wall antibiotics was driven specifically by a third (p)ppGpp synthetase, YwaC (SAS2). Without *ywaC*, cells are unable to form low-GTP persisters in response to treatment with cell wall antibiotics even in the presence of both *relA* and *yjbM* (Fig 6D). On the other hand, cells containing YwaC as the sole (p)ppGpp synthetase (r*elA^Syn^* Δ*yjbM*) retained the ability to induce low-GTP persisters in response to cell wall antibiotics (Fig 6D). This has a strong impact on cell survival. Deletion of *ywaC* not only modestly decreased survival upon sudden lethal treatment with cell wall antibiotics (Fig 6E); but more strikingly, completely abolished the adaptive survival to lethal antibiotics due to pretreatment with sublethal antibiotics (Fig 5D). These results collectively demonstrate the role of (p)ppGpp synthetase in antibiotic-induced, adaptive persister formation.

## Discussion

Persisters were proposed more than 70 years ago to describe a rare population of phenotypically switched cells that are refractory to killing by antibiotics (Bigger, 1944). Since then, great strides have been made to build the theoretical framework regarding the generation of persisters, such as the definition of triggered and spontaneous persisters, meanwhile outlining the challenges in defining antibiotic-induced persisters and characterizing molecular determinants underlying the phenotypic entrance into persistence (Balaban et al., 2019). Here we demonstrate, in the Gram-positive model bacterium *B. subtilis*, that multiple conditions lead to persister entrance by a core switch involving GTP depletion by (p)ppGpp, via different (p)ppGpp synthetases (Fig 7). We visualized, for the first time, spontaneous persister entrance in wild-type bacteria, demonstrating that (p)ppGpp is responsible for a rapid switch-like dynamics. We showed that persisters also form in response to bactericidal antibiotic treatment, even at sublethal concentrations, via (p)ppGpp-induction, revealing a pathway by which antibiotic treatment increases the reservoir of surviving cells for generation of antibiotic resistance. Since (p)ppGpp and its role in regulating purine synthesis is conserved in bacteria (Liu et al., 2015; Wang et al., 2018), persistence through (p)ppGpp-GTP antagonism is likely a widespread mechanism to allow survival against diverse antibiotics and other stresses.

**Figure 7.**
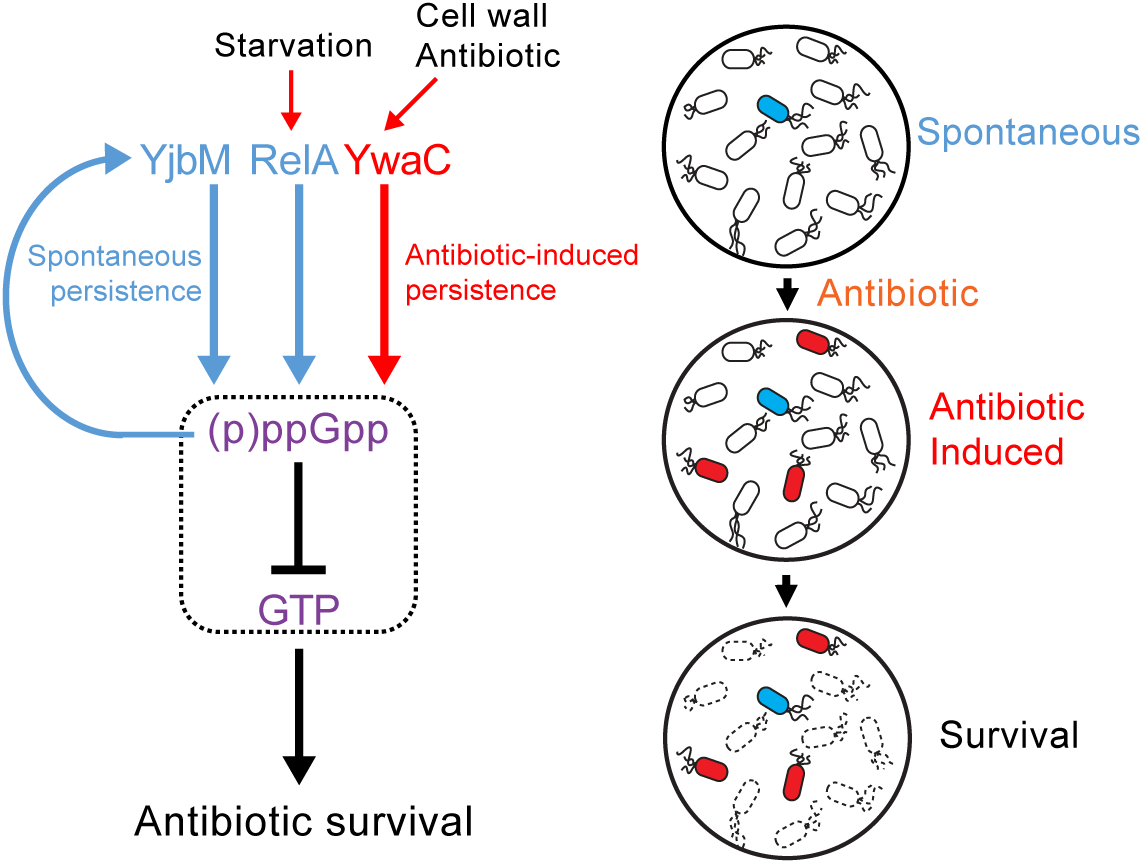
Model for persistence by (p)ppGpp-mediated GTP antagonism. Depletion of GTP is promoted by (p)ppGpp to enable multiple pathways to persistence. Induction of (p)ppGpp by starvation induces triggered persistence primarily through the (p)ppGpp synthetase RelA. Spontaneous persistence can occur during active growth through activities of two (p)ppGpp synthetases RelA and YjbM (spontaneous persistence, blue). Furthermore, persisters can also form in an antibiotic-responsive manner through YwaC (antibiotic-induced persistence, red). Thus, (p)ppGpp enables integration of different signals beyond starvation to effect persistence through GTP depletion. This allows multiple pathways of persister formation for antibiotic survival through a central metabolic switch.

### (p)ppGpp triggers the phenotypic switch to antibiotic persistence

Although (p)ppGpp has long been implicated in stationary phase or starvation-triggered persistence (Kaspy et al., 2013; Korch et al., 2003; Nguyen et al., 2011; Svenningsen et al., 2019), the causative role of (p)ppGpp in persistence has been unclear because starvation not only increases (p)ppGpp levels but also reduces ATP levels. The latter has been proposed to be the *bona fide* effector of persistence instead of (p)ppGpp (Conlon et al., 2016). Here our data demonstrate the critical role of (p)ppGpp in persister generation. First, we were able to purify rare high (p)ppGpp cells and show that they are almost all persisters (Fig 3E and 3F). Second, we confirmed that starvation-triggered persister formation is dependent on (p)ppGpp (Fig 1G). Thirdly, we found that ATP depletion, which was recently shown to trigger persistence (Conlon et al., 2016), also induces (p)ppGpp accumulation (Fig 1H). Fourth, we found that (p)ppGpp is necessary for switch-like spontaneous persister entrance dynamics (Fig 4G). Finally, we found that (p)ppGpp is also required for cell wall antibiotic-induced persistence (Fig 5). Importantly, genetic dissection of different (p)ppGpp synthetases revealed that they are differentially responsible for starvation-triggered, spontaneous, or antibiotic-induced persistence (Fig 6). Therefore, at least in *B. subtilis*, (p)ppGpp is a key inducer of persister generation and a modulator of persister dynamics. We suspect that (p)ppGpp also plays similar roles in persister generation in multiple pathogens, since mutants with elevated (p)ppGpp levels have been isolated from patients with recurrent infections after antibiotic treatments (Berti et al., 2018; Gao et al., 2010; Honsa et al., 2017).

There is a minor component (<10%) of (p)ppGpp-independent persistence due to sporulation (Fig 2D), a *B. subtilis* developmental pathway (Errington, 2003). Intriguingly, sporulation was shown to be promoted by (p)ppGpp indirectly through depletion of GTP (Vasantha and Freese, 1980). How the phenotypic switch and the developmental pathways crosstalk to protect cells in both short term and long period remain to be elucidated.

### GTP depletion is a key metabolic effector of persistence

We found that (p)ppGpp causes persistence by antagonizing GTP. Loss of persistence in the absence of (p)ppGpp can be completely compensated by a mutation that reduces GTP synthesis (Fig 2F). Furthermore, repression or disruption of purine biosynthesis genes alone is sufficient to cause a strong increase in persistence (Fig 2G and 2H). This suggests that although (p)ppGpp senses environmental signals to trigger the entrance to persistence, it is GTP depletion that is the *bona fide* effector of cell dormancy. This agrees with our previous observation that GTP is a metabolic gauge of growth rate and its depletion results in a suspension of cellular growth in *B. subtilis* (Bittner et al., 2014).

How does low GTP lead to growth suspension? Although reduction of GTP levels elicits a fundamental change in transcription (Krásný and Gourse, 2004; Kriel et al., 2014), the effect of depleting GTP is far beyond transcription. Low GTP is sufficient to broadly suspend diverse growth-determining cellular processes that utilize GTP, including DNA replication (Rymer et al., 2012), rRNA synthesis (Krásný and Gourse, 2004), translation initiation and elongation (Mitkevich et al., 2010), ribosome assembly (Britton, 2009), 100S ribosome dissociation (Tagami et al., 2012), and potentially cell wall synthesis (Bisson-Filho et al., 2017). In addition, (p)ppGpp may also compete with GTP for enzyme active sites (Corrigan et al., 2016; Kanjee et al., 2012) to exacerbate the inhibitory effect of low GTP. Since lethality of conventional antibiotics stems from disruption of these growth-associated processes (Kohanski et al., 2010), low GTP bacteria will be able to better survive antibiotic treatment. Therefore, persistence can be due to a global metabolic brake of multiple cellular processes triggered by low GTP. (p)ppGpp-GTP may also facilitate coordinated shut down of macromolecular synthesis, such as transcription and replication, to minimize detrimental conflicts (Liu et al., 2015). Finally, the growth suspension may also reduce generation of reactive oxygen species which have been implicated in antibiotic lethality (Dwyer et al., 2014). These alternative hypotheses require further investigation.

### Rapid switch in spontaneous persistence entrance via allosteric enzyme cooperativity

Spontaneous formation of persisters in unstressed populations is one of the most interesting examples of phenotypic switch (Veening et al., 2008). Here we were able to directly observe the single cell dynamics of spontaneous entrance to persistence in wild-type bacteria. We observe a striking switch-like dynamic, in which single cells elongate with normal speed and abruptly enter persistence. The entrance is via (p)ppGpp-induced depletion in GTP that reaches below a threshold (Fig S7B).

Phenotypic switch into persistence has been proposed to be due to the competition between toxins and antitoxins toxin-antitoxin (TA) which provides a threshold-like regulation (Fig S7A) (Balaban et al., 2004; Rotem et al., 2010). Our data suggest that cells can utilize a distinct alternative mechanism for the spontaneous persister entrance: through allosteric enzyme cooperativity and positive feedback in (p)ppGpp synthesis (Fig S7B). We showed that a mutation of allosteric pppGpp activation site in the synthetase YjbM resulted in significantly reduced spontaneous persistence to the same extent as the *yjbM* deletion mutant (Fig 6C). We propose that self-amplification of (p)ppGpp is required for spontaneous persister formation from a small input signal, such as gratuitous (p)ppGpp synthesis by YjbM or potentially RelA (Fig S7B). It was shown that RelA activity is also stimulated by pppGpp in *E. coli* (Kudrin et al., 2018). We hypothesize that this cooperativity may also contribute to spontaneous persister formation.

While (p)ppGpp accumulation in spontaneous persistence can be purely stochastic, we cannot rule out the possibility that (p)ppGpp could be triggered by transient stresses imposed on individual cells, such as localized nutrient fluctuation, or other stresses. For example, DNA damage has been shown to induce (p)ppGpp in *E. coli* (Kamarthapu et al., 2016). Whether these potential mechanisms are in play in Gram-positive bacteria remains to be explored.

### Antibiotic-induced persistence through the (p)ppGpp-GTP switch

Bigger’s original work not only suggest that persisters can be generated prior to antibiotic treatment, but also raised the possibility that persisters can be generated in response to antibiotics (Bigger, 1944). Subsequently, persisters were observed both before and after treatment with antibiotics (Balaban et al., 2019; Westfall et al., 2019). Our finding suggests that the persisters generated by exposure to antibiotics can at least in part be attributed to increases in the level of (p)ppGpp resulting from the stress imposed by these drugs. In addition, antibiotic-induced persisters can share the common metabolic signature with the persisters formed prior to antibiotic treatment: the (p)ppGpp-GTP dichotomy.

We found that treatment with cell wall antibiotics, either at lethal or sublethal concentration, promote persister formation by inducing accumulation of (p)ppGpp through the (p)ppGpp synthetase YwaC. *ywaC* is a component of the cell envelope stress regulon whose transcription is known to be induced by cell envelope damage (D’Elia et al., 2009; Eiamphungporn and Helmann, 2008; Libby et al., 2019), likely explaining its production of (p)ppGpp upon cell wall antibiotic stress. For the soil bacterium *B. subtilis*, the evolution of this regulation may be shaped by fitness advantage from its habitat to enhance survival to diffusible antimicrobials from neighboring microbes. For example, bacitracin, which can induce persister formation, is a diffusible natural antibiotic produced by other soil *Bacillus* species (Katz and Demain, 1977). This ppGpp-dependent mechanism of antibiotic-induced persistence is also likely conserved in pathogens occupying different niches. For example, the pathogen *S. aureus* has two different synthetases that can produce (p)ppGpp upon cell wall stress. Thus our model of drug-induced persisters may also explain why the (p)ppGpp mutants of *S. aureus* exhibit compromised survival against cell wall-damaging drugs (Geiger et al., 2014).

Antibiotic-induction of persistence by (p)ppGpp may promote the emergence of genetic resistance by increasing the reservoir of surviving bacteria under antibiotic assault. Even a brief exposure to sublethal dosage of bactericidal cell wall antibiotic can increase the surviving reservoir to up to 5-fold, allowing the evolution of antibiotic resistance in this population. Because human activities such as antibiotic abuse can lead to increased bacterial exposure to antibiotics which promotes the development of antibiotic resistance (Martinez, 2009), targeting persister formation may improve treatment efficacy and minimize the emergence of antibiotic-resistant bacteria.

### Author contribution

J.D.W. conceptualized the study and supervised the research. D.K.F. and J.L.T. designed the experiments. D.K.F., J.L.T. and D.Y. conducted experiments. J.W.S. performed bioinformatics analysis. D.K.F., J.L.T., J.W.S. and J.D.W. discussed the results and interpretations. D.K.F., J. L. T. and J.D.W. wrote the paper.

## Acknowledgements

We thank Alan Grossman, Kevin Griffith, and Dan Kearns for generously sharing strains and reagents. We thank Bruce Levin, Petra Levin, Michael Cox and members of the Wang lab for comments on the manuscript. This study is supported by HHMI Faculty Scholar Grant, NIH R35GM127088 and Hatch (to J.D.W).

## Materials and Methods

### Bacterial strains and strain construction

All bacterial strains, plasmids and oligonucleotides used in this study are listed in Table S1 to S3. LB and LB-agar were used for cloning and maintenance of strains. For selection in *B. subtilis*, media was supplemented with the following antibiotics when required: spectinomycin (80 µg/mL), chloramphenicol (5 µg/mL), kanamycin (10 µg/mL), and tetracycline (10 µg/mL). Combination of lincomycin (12.5 µg/mL) and erythromycin (0.5 µg/mL) was used to select for MLS resistance. Carbenicillin (100 µg/mL) was used for selection in *E. coli*. *B. subtilis* (p)ppGpp biosynthesis mutants were constructed by transformations of integration plasmids containing an I-sceI endonuclease cut site and regions of homology upstream and downstream of specific synthetase genes (pJW300 for Δ*ywaC*, and pJW371 for RelA^D264G^) followed by transformation of pSS4332 for Δ marker-less recombination (Janes and Stibitz, 2006). Successful recombination was verified by PCR. For the construction of (p)ppGpp^0^ mutant, Δ*ywaC* Δ*yjbM* was Δ*relA*::*mls* PCR product from genomic DNA using oligos oJW418/oJW419 and selected for MLS resistance (Kriel et al., 2012).

Construction of integration plasmid for YjbM^F42A^ was done by site-directed mutagenesis of pJW370 by PCR using oligos oJW2309/oJW2310.The *gmk*^Q110R^ allele was generated from a suppressor screen on guanosine-containing plates (Kriel et al., 2012). *B. subtilis* deletion mutants were constructed by serial transformation of PCR products from the *B. subtilis* knockout collection (BGSC, Gross lab). Where appropriate, Cre recombinase-mediated removal of the lox-site flanked *erm*^R^ cassette with pDR244-*cre* was performed followed by selection for the loss of MLS resistance. Construction of fluorescence reporters was done by fusion of PCR products containing respective promoter regions (primers oJW1935/oJW1936 for P*_ilvB_* or oJW2083/oJW2084 for P*_rrnB_*) with coding regions of fluorescence proteins (primers oJW1995/oJW1996 for GFP or oJW2805/oJW2806 for mCherry) using ligase cycling reaction (LCR) (de Kok et al., 2014). In the case of P*_ilvB_*-GFP or P*_ilvB_*-mCherry fusions, the construct was cloned into pDR110 flanked by *amyE.* In the case of P*_rrnB_*-GFPns (unstable GFP sequence from Griffith *et al*. (Griffith and Grossman, 2008)), DNA fragments of GFPns (primers oJW1995/oJW2020), flanking regions of *lacA* (primers oJW1990/oJW2414 and oJW2413/oJW2082), and lox-site flanked *erm*^R^ cassette (primers oJW2133/oJW2134) were amplified by PCR. The resulting PCR products were fused by LCR followed by amplification using PCR to generate linear recombination fragment of *lacA*::P*_rrnB_*-GFPns. Same method was used to construct *lacA*::P*_ywaC_*-mCh using oligonucleotide fragments of P*_ywaC_* (primers oJW3099/oJW3079). Chromosome integration of reporter constructs was done by transformation and selection for SPEC or MLS resistance. Removal of the lox-site flanked *erm*^R^ cassette with was done by transformation with pDR244-*cre* and selected for the loss of MLS resistance. All mutants and constructs were verified by DNA sequencing.

The (p)ppGpp^0^ *gmk*^Q110R^ mutant was obtained from isolating suppressor mutants from (p)ppGpp^0^ cells by plating on S7 minimal medium plate containing 1% glucose. The surviving colonies were plated on S7 minimal medium plate containing 0.5% casamino acids and 0.5 mM 8-azaguanine, or S7 minimal medium plate containing 0.5% casamino acids and 0.1 mM guanosine (Kriel et al., 2012) to differentiate between mutants containing mutation in HprT or Gmk. Colonies which can grow on guanosine but not 8-azaguanine were sequenced to identify the mutant *gmk* allele.

For strains that contained deletions of resident prophages. Whole genome sequencing was performed after genetic manipulations to ensure no re-introduction of prophages occurred.

### Growth conditions

*Bacillus subtilis* strains were grown in S7 defined medium (Vasantha and Freese, 1980); MOPS was used at 50 mM rather than 100 mM, supplemented with 0.1% glutamate, 1% glucose, and 0.5% casamino acids. Growth of YB886 strain background was supplemented with 20 µg/mL tryptophan and 50 µg/mL methionine (Harwood and Cutting, 1990). Cell cultures were grown at 37 °C from overnight plates with vigorous shaking. Cultures in logarithmic phase (OD_600_ = 0.1-0.3) were treated with antibiotics or inducers, such as arginine hydroxamate (RHX, 0.5 mg/mL), carbonyl cyanide m-chlorophenyl hydrazone (CCCP, 5 µM), sodium azide (NaN_3_, 4 mM), or arsenate (2.5-D-1-thiogalactopyranoside (IPTG) was added to a final concentration of 0.5 mM to induce *guaB* expression from an IPTG-inducible promoter (P*_spac_*). For microscopy experiments, the following inducers and concentrations were used-methylglucoside (-MG): 5%; CCCP: 5 unless otherwise specified: RHX: 0.5 mg/mL; μg/mL (0.5x MIC) or 100 μg/mL (2x MIC); CIP: 0.1 μg/mL (4x MIC); VAN: 0.1μg/mL (20x MIC)) along with non-induction controls. For nutrient downshift experiments, cells grown in S7 media supplemented with casamino acids were washed three times with S7 minimal media and re-suspended in an equal volume of S7 minimal media followed by 30-min incubation under the same condition.

### Minimum inhibitory concentration (MIC) determination

MICs for chloramphenicol (CAM), tetracycline (TET), kanamycin (KAN), ciprofloxacin (CIP), norfloxacin (NOR), rifampicin (RIF), bacitracin (BAC) and vancomycin (VAN) were determined using the microdilution method (Adimpong et al., 2012). Logarithmic phase cells were back-diluted to a final titer of ≈ 5×10 CFU/mL into 96-well plates containing two-fold serial dilutions of respective antibiotics in S7 defined medium. After 16-20 hours of incubation at 37 °C with shaking, the MIC was determined as the lowest drug concentration that prevented visible growth.

### Persister assay

Cells were harvested from young, overnight LB-agar plates (< 12 h), back-diluted into fresh S7 defined media at OD_600_ = 0.005, and grown at 37 °C with vigorous shaking. Cells were grown to logarithmic phase (OD_600_ ≈ 0.1-0.3). Treatments with bactericidal μg/mL; VAN, 4 μg/mL. To determine cell viability, culture aliquots were taken at T=0 and at designated times after treatment, plated onto Luria-Broth (LB) agar, and incubated at 37°C overnight. Viability at different time points was determined as CFU/mL/OD_600_ and relative survival (vs T_0_) was calculated. For experiments involving pre-induction of cells with (p)ppGpp-inducing agents, cells were grown to OD_600_ ≈ 0.1 and divided into two: g/mL; CCCP, 5 μ NaN_3_, 4 mM; arsenate, 2.5 mM) and other as non-induction control. The cultures were grown for an additional 30 min under the same conditions (T=0.5 hr) and subjected to the persister assay, as described above. In the case of BAC treatment, the final antibiotic concentrations were identical for both induced and non-induced populations.

### Measurement of intracellular nucleotides by thin-layer chromatography

To measure intracellular nucleotides, cells were first harvested from overnight plates, back-diluted to OD_600_ = 0.005, and grown in low-phosphate S7 defined medium (0.1X phosphate, 0.5 mM), supplemented with casamino acids. Once cultures reached OD_600_ ≈ 0.05, 1 mL cells were labeled with 50 µCi of ^32^P orthophosphate (900 mCi/mmol; Perkin Elmer) or 2-3 generations before treatment or sampling. At OD_600_ ≈ 0.15, RHX, CCCP, or arsenate were added to the cultures and samples were collected at regular time points for nucleotide extraction. Nucleotides were extracted by incubating 100 µL cells with 20 µL of 2 N formic acid for at least 20 minutes. Samples were spotted on PEI cellulose thin-layer chromatography (TLC) plates (Selecto) and developed in 1.5 M or 0.85 M potassium phosphate monobasic (KH_2_PO_4_, pH 3.4) buffer to separate (p)ppGpp or GTP, respectively. TLC plates were exposed on storage phosphor screens (GE Healthcare) and scanned on a Typhoon imager (GE Healthcare).

### Fluorescence plate assay

To monitor GFP expression from P*_ilvB_*-GFP reporter during growth in liquid culture, cells were grown to OD_600_ ≈ 0.1-0.3 in S7 defined medium and then diluted to OD_600_ = 0.005 into fresh S7 defined medium. 200 µL of the diluted cultures were transferred into wells of 96-well black plate with flat clear bottom and transparent lid. The plate was incubated at 37°C with constant shaking for up to 18 h in a Biotek Synergy2 microplate reader. Measurement of OD_600_ and GFP signals of individual wells was performed every 10 min using absorbance monochromator (for OD_600_), and excitation (485/20) and emission filters (528/20) for GFP.

### Fluorescence microscopy

To monitor (p)ppGpp induction using fluorescence reporters, cells were grown to OD_600_ ≈ 0.1 followed by 30-min induction with RHX, α CARB, BAC, CIP, KAN, or VAN (concentrations were listed under growth conditions) along with non-induction controls. All imaging samples were spotted on 1.5% agarose pads made with the same growth medium, and immediately imaged with Olympus IX-83 inverted microscope (Olympus) using 60X phase contrast objective with fluorescence filters (excitation: 470/20 nm, dichroic mirror: 485 nm, emission: 515/50 nm for GFP; excitation: 575/20 nm, dichroic mirror: 595 nm, emission: 645/90 nm for mCherry; and excitation: 427/10 nm, dichroic mirror: 595 nm, emission: 472/30 nm for Sytox blue). Single-cell time-lapse imaging was performed at 37°C using temperature-controlled imaging chamber (Tokai Hit) coupled with automatic stage and microscope control as described previously (Young et al., 2011). For imaging persister survival, 5 μ and 25 nM Sytox blue (Molecular Probes) at final concentrations were used. Strains without the fluorescence reporters were used for autofluorescence measurement.

### Flow cytometry and cell sorting

Flow cytometry was performed in UWCCC flow cytometry core. Cells for flow cytometry analysis were grown under same experimental conditions as described above. For flow cytometry analysis, cells were immediately fixed with 0.4% paraformaldehyde for 15 min at room temperature, washed 3x with 1x phosphate buffered saline (PBS), and kept at 4°C until analysis. Fixation was verified by viability plating and microscopy. Flow cytometry analysis was performed using BD LSRFortessa flow cytometer (BD Biosciences) with a 70-µm nozzle. Cell populations were detected using both forward and side scatter (FSC and SSC). Single-cell fluorescence was measured using 488 nm laser and detection filters for GFP (530/30 nm, 505LP dichroic filter). Approximately two million events were counted for each sample and. FACS analysis was performed using BD FACSAria cell sorter (BD Biosciences) with a 70-µm nozzle at room temperature using 488 nm laser and detection filters for GFP (525/50 nm, 505LP dichroic filter). At least 1,000 cells were obtained from the rarest gate for each sample. Cell recovery rate was estimated to be > 90% based on viability counting on LB plates. Aliquots of gated populations were subjected to persister assay with 4X MIC of VAN as described above. We confirmed that this antibiotic dosage does not affect the killing (Fig 1B) but facilitate drug removal for plating from sorted fractions with low cell numbers.

### Transposon sequencing (Tn-Seq)

*B. subtilis* 168 transposon mutant library was kindly provided by the Grossman Lab (Johnson and Grossman, 2014). Construction of the library is briefly described as follows: *In vitro* transposition of B. subtilis 168 genomic DNA (gDNA) with magellen6x transposon was performed by mixing 1.3 µg pCJ41 (containing magellen6x transposon), 34 ng purified MarC9 transposase, 5 µg B. subtilis gDNA, 10 µL 2x buffer A (41 mM HEPES pH 7.9, 19% glycerol, 187 mM NaCl, 19 mM MgCl2, 476 µg/mL BSA and 3.8 mM DTT) into 20 µL reaction *in vitro* and incubated overnight at 30°C. The transposed DNA was precipitated and resuspended in 2 µL 10x buffer B (500 mM Tris-Cl pH 7.8, 100 mM MgCl2, 10 mM DTT), 2 µL 1 mg/mL BSA and 11 µL H2O followed by 4h incubation at 37°C. After incubation, 4 µL 2.5 mM dNTPs and 1 µL of 3U/ µL T4 DNA Polymerase were added to the DNA and further incubated for 20 min at 12°C, followed by heat inactivation at 75°C for 15 min. Next, 0.2 µL 2.6 mM NAD and 1 µL of 10 U/ µL *E. coli* DNA ligase were added and the reaction was incubated at 16°C overnight. The resulting in vitro transposed and repaired gDNA was transformed into *B. subtilis* 168 and plated on LB agar containing spectinomycin and incubated overnight. Colonies containing the transposon were washed off and pooled into a single library. The library was estimated to contain ∼50,000 unique transposon inserts across the genome.

For the selection experiment with the transposon library, an aliquot of the library was inoculated and grown in S7 medium supplemented with tryptophan (20 µg/mL) to OD_600_ ≈ 0.1-0.3 and subjected to VAN or CIP treatment as described above. Survivors before and after antibiotic treatment were plated onto LB plates and recovered after ∼14 h incubation at 37°C. ∼650,000 colonies from each sample were pooled and froze down for genomic DNA extraction and sequencing library preparation.

Preparation of sequencing library was performed as previously described (van Opijnen et al., 2014). Frozen cell pellets were resuspended in 500 µL lysis buffer with lysozyme and RNAse A (20 mM Tris-HCl pH 7.5, 50 mM EDTA, 100 mM NaCl, 2 mg/mL lysozyme, 120 µg/mL RNAse A) and incubated at 37°C for 20-30 min. Next, the incubated cell lysate was mixed with 60 uL 10% N-lauroylsarkosine and further incubated at 37°C for 15 min. Genomic DNA (gDNA) was purified using 600 µL phenol, then 600 µL phenol:chloroform:isoamyl alcohol (25:24:1), and finally 600 µL pure chloroform. DNA in the aqueous phase was precipitated using 1/10 volumes of 3M NaOAc and 2 volumes of 100% ethanol. DNA pellet was washed with 70% ethanol, air-dried on bench, and resuspended in 10 mM Tris-HCl, pH 8.5 and stored at 4°C. For each sample, 6 µg of DNA was used for MmeI digestion in 200 µL (6 µg gDNA, 6 µL MmeI (2000 U/mL, NEB), 0.5 µL 32 mM S-adenosylmethionine, 20 µL NEB CutSmart Buffer, and ddH_2_O up to 200 µL). DNA was digested for 2.5 h at 37°C, after which 2 µL calf intestinal phosphatase (10,000 U/mL, NEB) was added to the digest and incubated for 1h at 37°C. Digested gDNA was extracted with 200 µL phenol:chloroform:isoamyl alcohol (25:24:1) followed by 200 µL of pure chloroform. DNA in the aqueous phase was first mixed with 1/10 volume of 3M NaOAc and 67 ng/mL glycogen, and then 2.5 volumes of 100% ethanol. The tubes were then placed at −80°C for 20 min and then centrifuged at max speed for 15 min at 4°C. Precipitated DNA was washed with 150 µL 70% ethanol twice at room temperature, air-dried, and resuspended in 15 µL ddH_2_O.

For annealing of DNA adaptor, 20 µL of 100 µM synthesized oligos (IDT) were mixed with 1 µL of 41 mM Tris-HCl pH 8.0 (final concentration of adaptor: 50 µM in 1mM Tris-HCl pH 8.0). Oligos were annealed by heat denaturation (95 °C for 5 min) and stepwise cool-down (94°C for 45 s then repeat with −0.3°C per cycle for 250 cycles, then hold at 15°C) using PCR machine. Annealed adaptors were diluted to 3.3 µM in ddH2O and stored at −20°C. For adaptor ligation, 5 µL of digested DNA was mixed with 1 µL of 3.3 µM DNA adaptor, 1 µL of 10x T4 DNA ligase buffer (NEB), 1 µL of T4 DNA ligase (400,000U/mL, NEB) and 2 µL ddH2O. The ligation mix was incubated overnight at 16°C in a PCR machine.

Amplification of adaptor-ligated DNA library was performed using barcoded primers and Phusion high-fidelity DNA polymerase (NEB) for 18 cycles according to provided instructions. The PCR products were mixed in equal amounts, purified by size-exclusion, and submitted for sequencing using Illumina sequencing primer (5’-ACACTCTTTCCCTACACGACGCTCTTCCGATCT-3’). Deep sequencing was done using Illumina HiSeq 2500 (Illumina) by the University of Michigan DNA Sequencing Core. Analysis of sequencing data was done using a custom Python script and mapped to the *B. subtilis* 168 reference genome (NC_000964.3). Visual inspection of transposon insertion profiles was done using GenomeBrowse (Golden Helix).

### Quantification and Statistical Analysis

For TLC experiments, intensities of nucleotide spots were quantified using ImageQuant software (Molecular Dynamics) and normalized to the number of phosphates in the corresponding nucleotide and ATP level at T=0 for comparison between samples.

Microscopy image analysis and cell parameters (cell area and fluorescence intensity) measurements were done using Metamorph software (Molecular Devices). Single cell growth rate (µ) at each frame was calculated using the equation: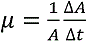
, where A is t is the change in time (in min). Flow-cytometry data were analyzed using FlowJo X software (FlowJo, LLC). Smallest allowed gate was used for forward and side scatter to reduce variations in fluorescence signals due to size variation. Identical gating strategies were used for different samples.

Statistical information of individual experiments are included in the figure legends. n represents the number of biological replicates or number of cells for experiments involving single-cell measurements as indicated. Significance was tested using Student’s t test. In figures, asterisks indicate statistical significance between datasets where *p < 0.05, **p < 0.005, ns, not significant. Prism 7 (GraphPad) was used for statistical analysis.

---

**Figure S1.**
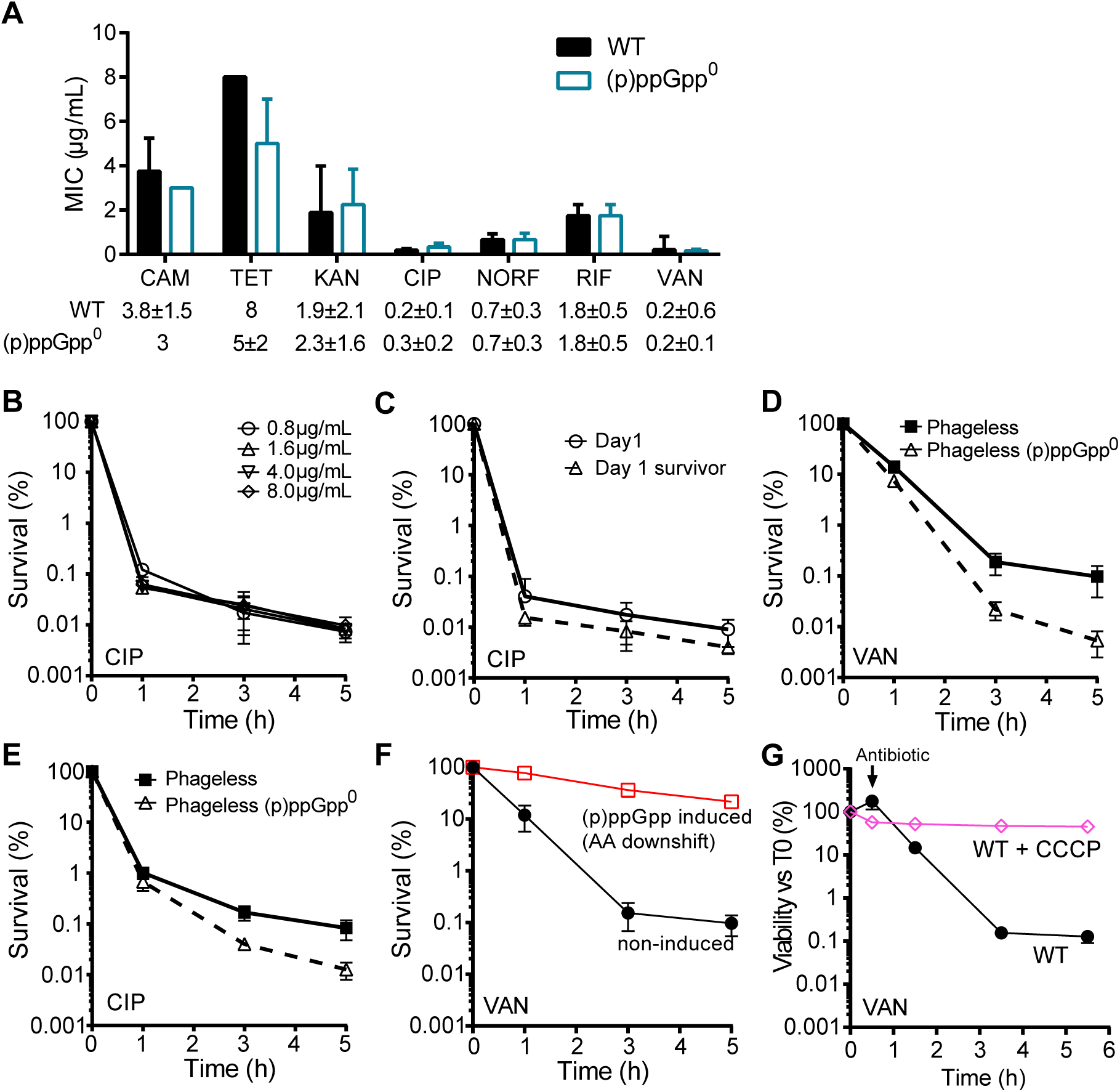
(p)ppGpp mediated persistence is not due to changes in drug sensitivities or prophage contamination. (A) Minimal inhibitory concentrations (MICs) of WT and (p)ppGpp^0^ to various antibiotics. (B) Survival of WT cells to concentrations of CIP ranging from 4x to 40x MIC for up to 5 h. (C) Survival of exponentially-growing WT cells (Day 1, solid lines), or its successor population derived from survivors after 5 h treatment (Day 1 survivor, dashed lines) to CIP for up to 5 h. (D and E) Survival of WT and (p)ppGpp^0^ mutant in the prophage-cured background (Δ*zpdN* ΔSP β ΔPBSX) to (D) VAN or (E) CIP treatment. (F) Survival of (p)ppGpp-induced and non-induced WT cells to VAN. (p)ppGpp induction was performed by growth transition from rich media to amino acids depleted media for 30 min. (G) WT cells with or without 30 min treatment of ATP synthesis inhibitor CCCP were tested for persister levels with VAN treatment for 5 h. All data points represent mean ± SD, n = 3.

**Figure S2.**
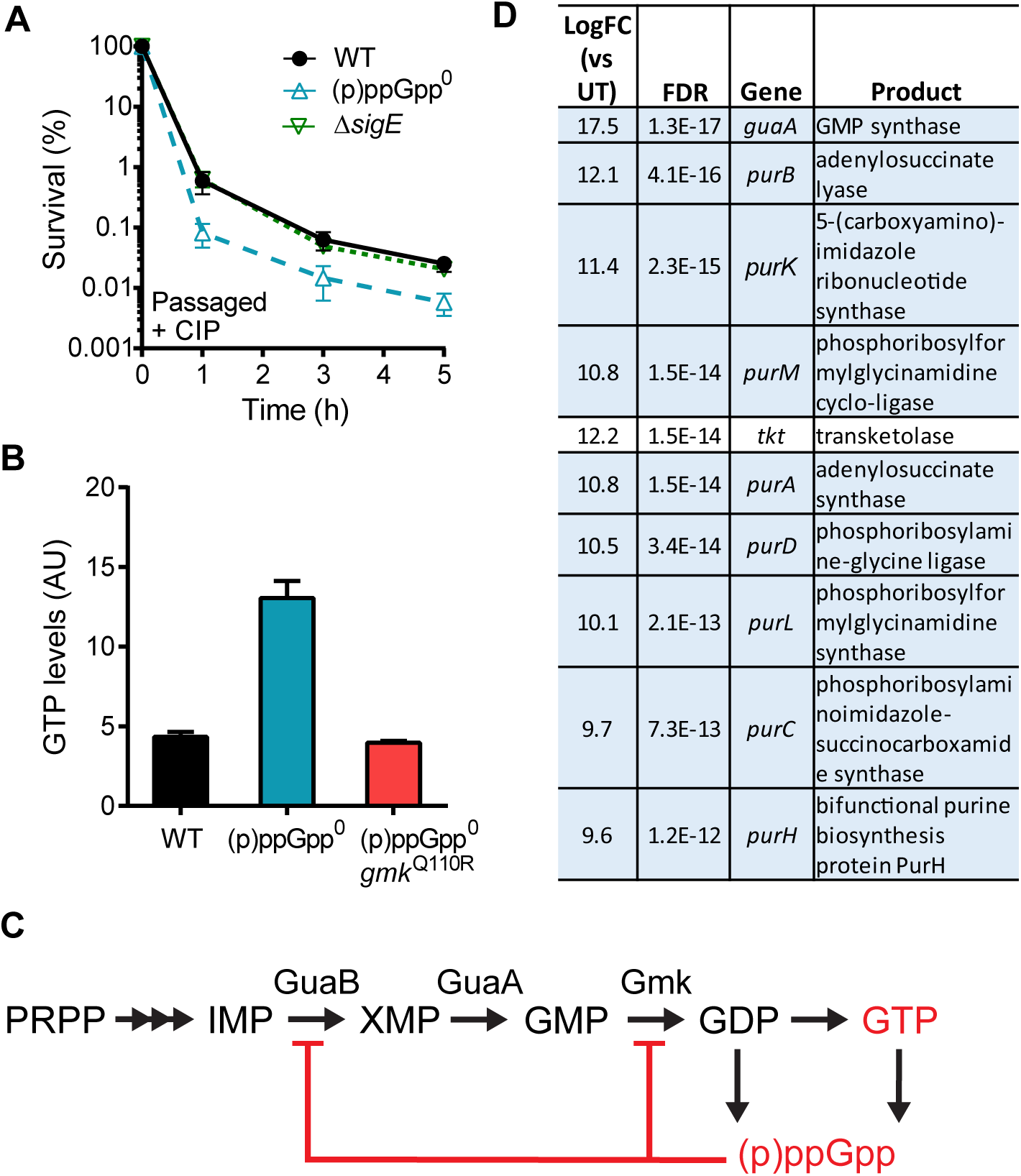
Persistence by GTP depletion. (A) Spontaneous persistence is unaffected by loss of sporulation. Survival of passaged and exponentially-growing WT, (p)ppGpp^0^, and Δ*sigE* cells to CIP treatment over time. Values represent mean ± SD, n = 2. (B) GTP levels of WT (black), (p)ppGpp^0^ (blue), and (p)ppGpp^0^ *gmk*^Q110R^ (red). Values represent mean ± SD, n = 3. (C) Schematic of *de novo* purine and downstream GTP biosynthesis. The *pur* operon encodes enzymes responsible for the conversion of PRPP to IMP, which leads to downstream GTP biosynthesis. Gmk and GuaB are directly inhibited by (p)ppGpp as indicated. (D) Tables showing the top ten genes that are enriched for disruptions after CIP treatment. LogFC represents fold enrichment in Log2. Genes highlighted in blue are involved in purine biosynthesis. FDR represents false discovery rate.

**Figure S3.**
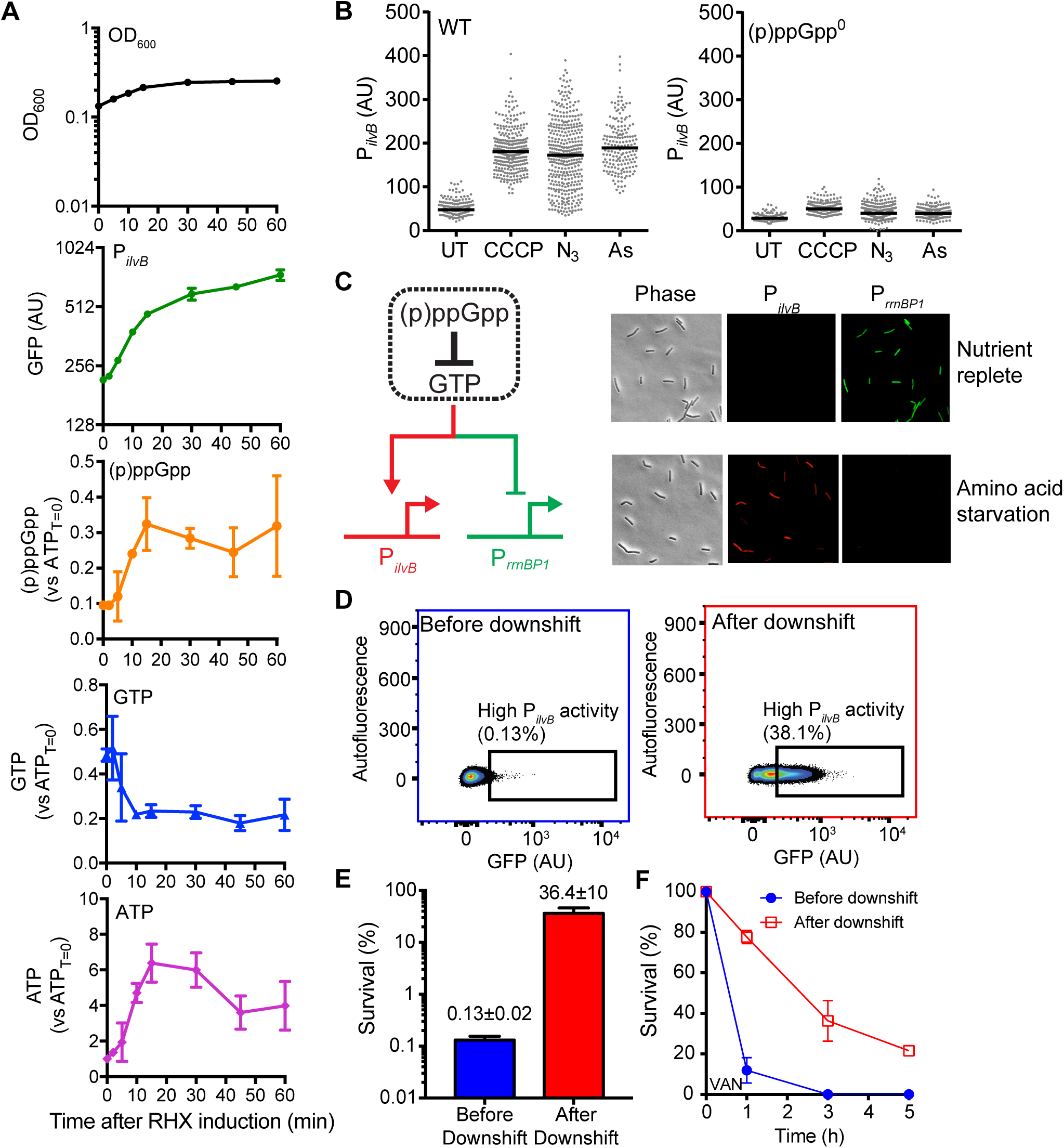
P*_ilvB_* activities to chemical and physiological induction of (p)ppGpp. (A) Time-dependent response of P*_ilvB_* activities and nucleotide levels to amino acid starvation. WT cells were grown to log phase and treated with 0.5 mg/mL RHX followed by measurement of growth (OD_600_), P*_ilvB_* activities (GFP) and relative abundance of various intracellular nucleotides as indicated. Fluorescence was measured using fluorescence microscopy and nucleotide levels were determined using TLC. Each time point represents mean ± SD, n = 3. (B) P*_ilvB_* activities of WT and (p)ppGpp cells exposed to 5 µM0 CCCP, 2.5 mM arsenate or 4 mM sodium azide for 30 min. UT represents untreated control. (C) Depletion of GTP increases the activity of *ilvB* promoter (i.e. expression of P*_ilvB_*-*mcherry*) but inhibits the ribosomal P1 promoter (i.e. inhibition of P-*gfp*^ns^ expression). *gfp*^ns^ encodes a degradable variant of GFP (see Methods). Representative images shown are WT cells containing both reporters before and after amino acids starvation. (D to F) Amino acid starvation increases the level of low GTP persisters. (D) Flow-cytometry analysis of WT populations before and after amino acids downshift using the P*_ilvB_* reporter. Fraction of GFP-high cells above the threshold in the population was measured based on FACS sorting analysis in Fig 3F. (E and F) Survival of populations from (D) to VAN treatment for (E) 3 h or (F) over a period of 5 h. Values represent mean ± SD, n > 10^6^ cells each, 3 replicates.

**Figure S4.**
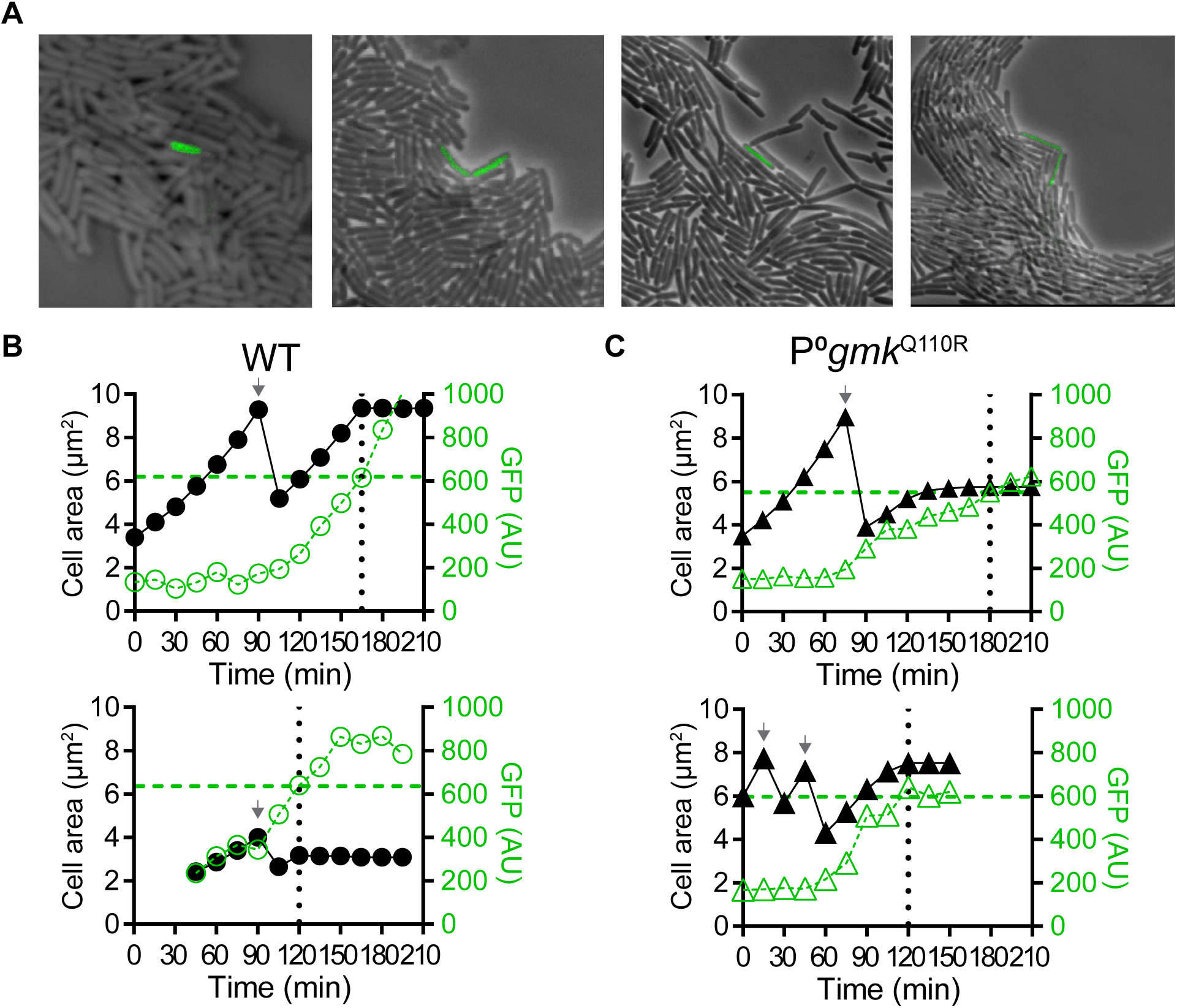
related to Fig. 5. Spontaneous development of GTP-depleted persisters. (A) Snapshot images of spontaneous persisters in WT cells containing the P*_ilvB_* reporter. (B and C) Independent events of persistence entrance in WT and (p)ppGpp^0^ *gmk*^Q110R^ mutant in addition to data shown in Fig 4E. Changes in P*_ilvB_*-*gfp* (green) and cell size (black) were shown during transition into persistence in (B) WT and (C) (p)ppGpp^0^ *gmk*^Q110R^ mutant. Persistence entrance was marked by vertical dotted lines indicating time and horizontal dashed line indicating P*_ilvB_*-*gfp*. Arrows indicate cell division.

**Figure S5.**
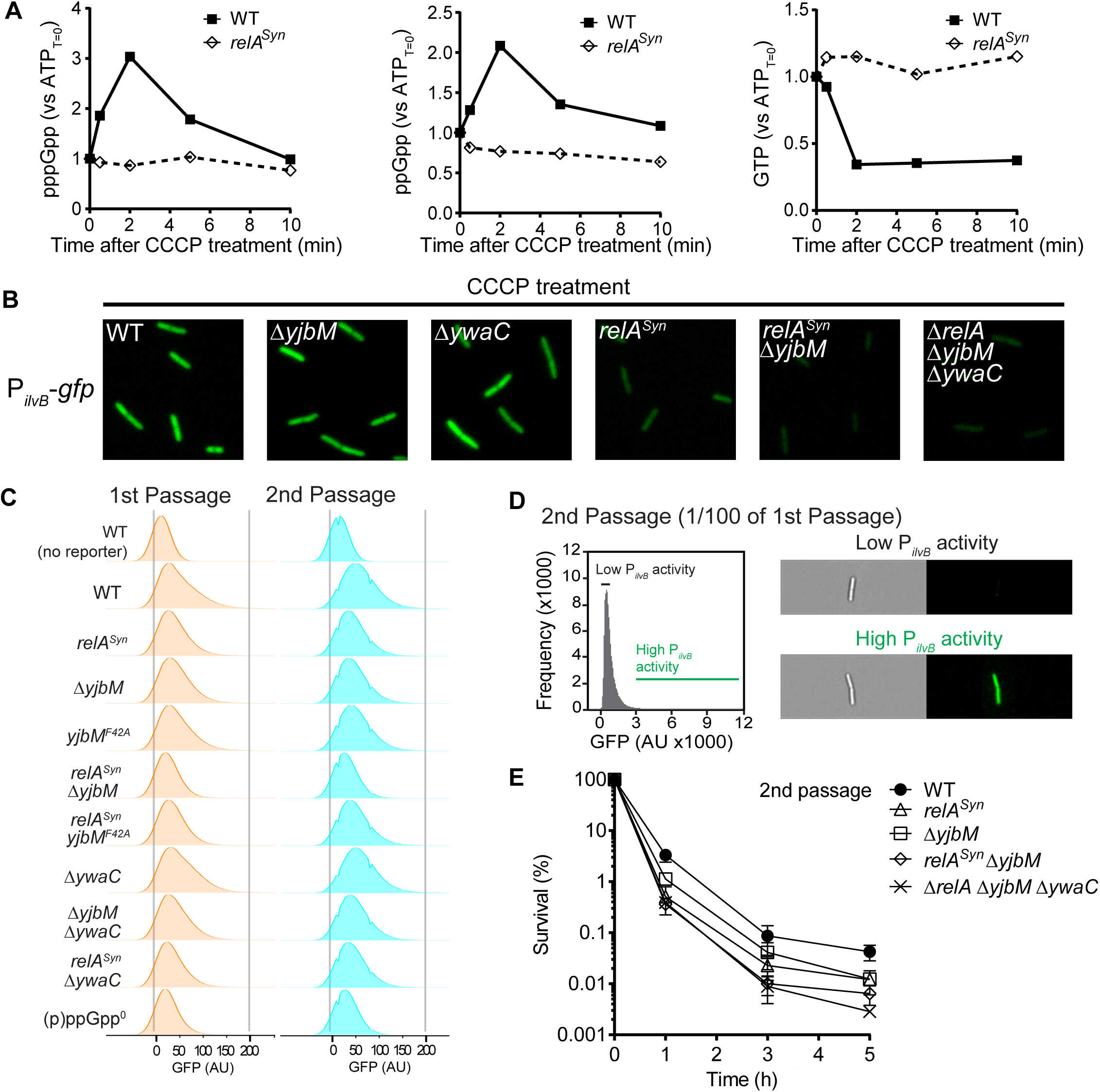
Role of different (p)ppGpp synthetases on (p)ppGpp synthesis and persistence. (A) Inhibitor of ATP synthesis induces (p)ppGpp through RelA. WT and relA^Syn^ cells were treated with CCCP and measured for changes in pppGpp, ppGpp, and GTP with TLC over time as indicated. (B) Snapshots of WT and (p)ppGpp synthetase mutants containing the P_ilvB_ reporter treated with CCCP for 30 min. (C) Flow cytometry analysis of wild-type and (p)ppGpp biosynthesis mutants containing the PilvB reporter (∼1.5×10^6^ cells each). Cells were grown to exponential phase (1st Passage, orange), diluted by 100-fold into fresh media and re-grew to exponential phase (2nd Passage, cyan). Both generations were analyzed by flow cytometry to measure the level of low-GTP cells. See also Figure S6. Values represent mean ± SD from three independent replicates of n > 10^6^ cells each. (D) Representative images of cells with low or high P_ilvB_ activity in a passaged population of exponentially growing WT cells. (E) Survival of WT, *relA*^Syn^, Δ*yjbM*, *relA*^Syn^Δ*yjbM* and Δ*yjbM*Δ*ywaC*Δ*relA* to VAN treatment over time. Cells were grown to and passaged at exponential phase before treatment. Values represent mean ± SEM, n = 3.

**Figure S6.**
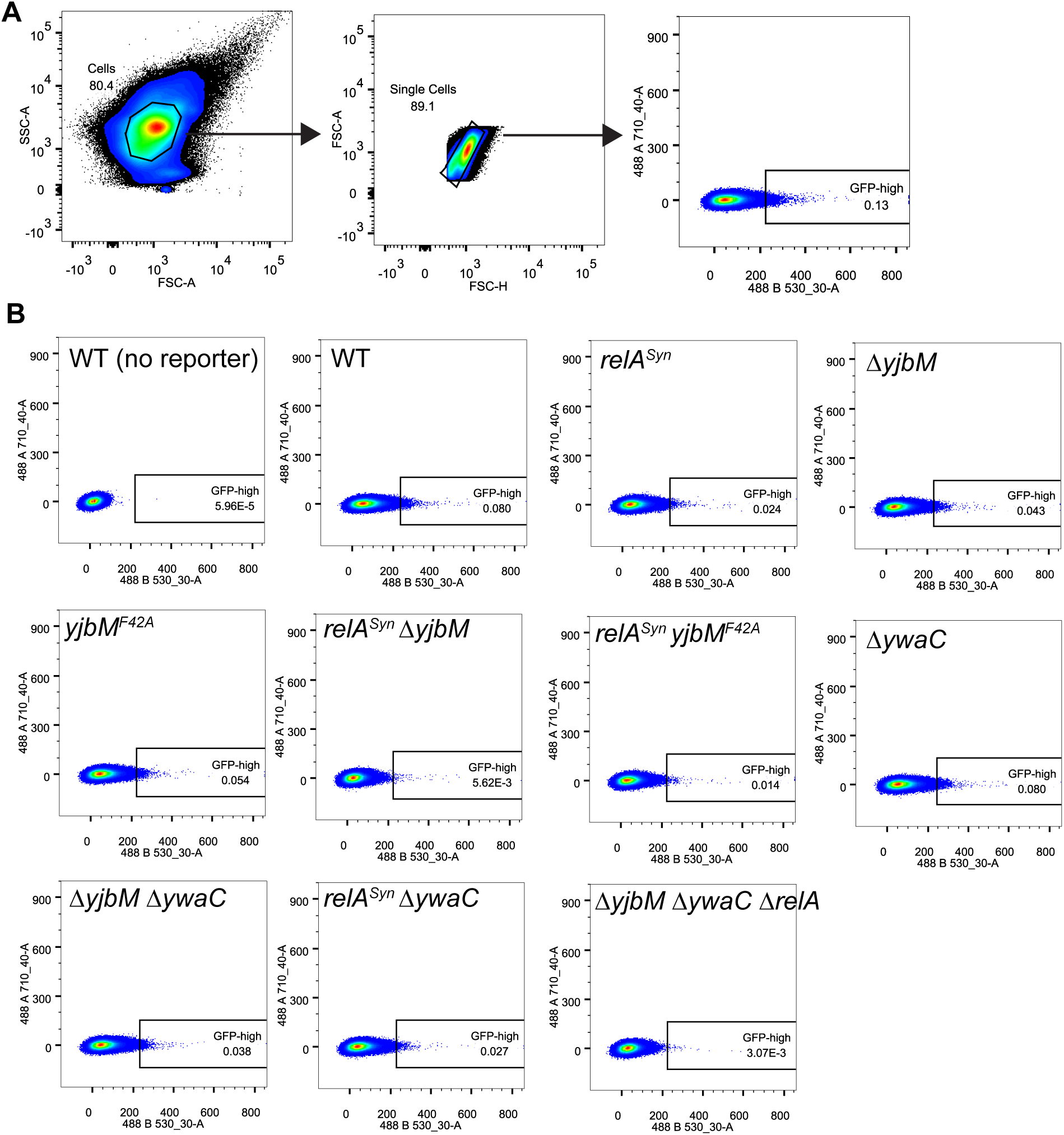
Flow cytometry analysis of low-GTP persister frequencies. (A) Gating strategy for measurement of cells with high P_ilvB_ activity. Cells within a narrow range of cell sizes (”Cells”) were gated, sub-gated to filter cell aggregates (”Single cells”), and analyzed for their fluorescence distribution. A fluorescence cut-off determined using wild type by our FACS experiment (∼3-fold above mean fluorescence) was used to determine the level of cells with high P_ilvB_ activity. The same cut-off was used for both wild type and (p)ppGpp synthetase mutants. SSC: side-scatter, FSC: forward-scatter, 488 A710_40: autofluorescence, 488 B530_30: GFP fluorescence. (B) Representative scatter plots of passaged growing populations of wild type and (p)ppGpp synthetase mutants. Numbers indicate the fraction (%) of GFP-high cells. ∼1.5×10^6^ cells for each.

**Figure S7.**
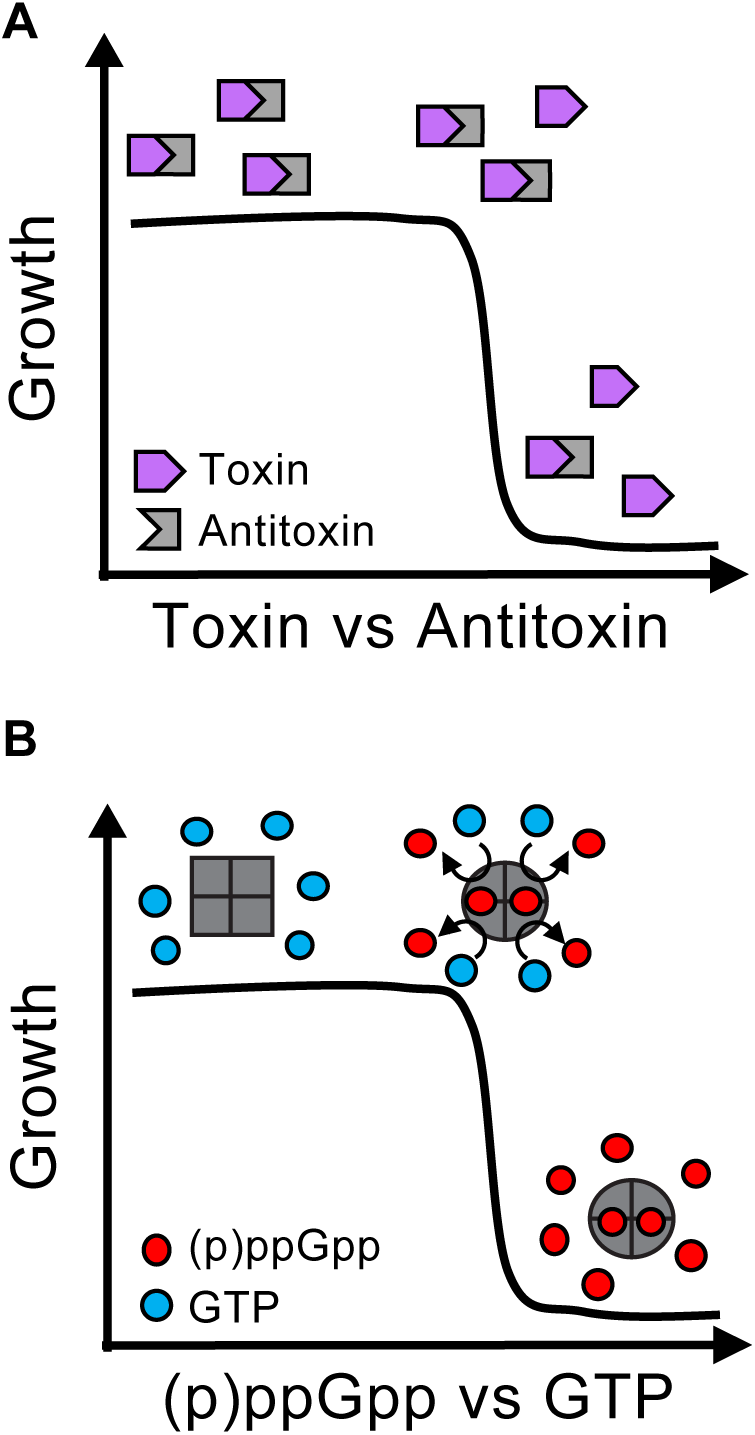
Switch-like persister formation by toxin-antitoxin competition or enzyme cooperativity. (A) Persistence regulation by toxin-antitoxin (TA) systems. The switch between growth and persistence is regulated by the relative abundance of toxins or antitoxins. Cells become dormant persisters when toxins exceed antitoxins by a threshold level. (B) Persistence control by enzyme cooperativity. The phenotypic switch into persisters is determined by allosteric activation of oligomeric (p)ppGpp synthetases (e.g. YjbM) by pppGpp. The self-amplification of (p)ppGpp synthesis leads to rapid depletion of GTP below a critical threshold leading to dormancy.

**Table S1.** Bacterial strains used in this study

|  | Genotype | Source |
| --- | --- | --- |
| JDW2144 | 3610 *coml*<sup>Q21L</sup> Prototroph | Daniel Kearns |
| JDW2231 | JDW2144 $\Delta yjbM \Delta ywaC \Delta relA::mIs$ | This work |
| JDW2234 | JDW2144 $\Delta pyk$ | Lab collection |
| JDW2528 | JDW2144 $\Delta yjbM \Delta ywaC \Delta relA gmk^{Q110R}$ | This work |
| JDW2721 | JDW2144 *relA*<sup>D264G</sup> | This work |
| JDW2901 | 3610 $\Delta zpdN \Delta SP\beta \Delta PBSX \Delta comI$ | Daniel Kearns |
| JDW2911 | JDW2144 $\Delta ydcE::erm \Delta yonT \Delta txpA$ | This work |
| JDW2914 | JDW2144 $\Delta codY$ | This work |
| JDW2921 | JDW2144 *amyE::P<sub>ilvB</sub>-GFPmut2*<sup>A206K</sup> | This work |
| JDW2923 | JDW2144 $\Delta yjbM amyE::P_{ilvB}\text{-GFPmut2}^{A206K}$ | This work |
| JDW2927 | JDW2144 $\Delta yjbM \Delta ywaC \Delta relA amyE::P_{ilvB}\text{-GFPmut2}^{A206K}$ | This work |
| JDW2929 | JDW2144 *comK::erm* | This work |
| JDW2935 | JDW2144 *relA*<sup>D264G</sup> *amyE::P<sub>ilvB</sub>-GFPmut2*<sup>A206K</sup> | This work |
| JDW2937 | JDW2144 $\Delta ywaC amyE::P_{ilvB}\text{-GFPmut2}^{A206K}$ | This work |
| JDW2939 | JDW2144 $\Delta yjbM \Delta ywaC amyE::P_{ilvB}\text{-GFPmut2}^{A206K}$ | This work |
| JDW2941 | JDW2144 *relA*<sup>D264G</sup> $\Delta yjbM amyE::P_{ilvB}\text{-GFPmut2}^{A206K}$ | This work |
| JDW2947 | JDW2144 $\Delta ywaC relA^{D264G} amyE::P_{ilvB}\text{-GFPmut2}^{A206K}$ | This work |
| JDW2949 | JDW2144 *yjbM*<sup>F42A</sup> *amyE::P<sub>ilvB</sub>-GFPmut2*<sup>A206K</sup> | This work |
| JDW2951 | JDW2144 *yjbM*<sup>F42A</sup> *relA*<sup>D264G</sup> *amyE::P<sub>ilvB</sub>-GFPmut2*<sup>A206K</sup> | This work |
| JDW2953 | JDW2144 $\Delta yjbM \Delta ywaC \Delta relA gmk^{Q110R} amyE::P_{ilvB}\text{-GFPmut2}^{A206K}$ | This work |
| JDW2961 | JDW2144 *amyE::P<sub>ilvB</sub>-mCherry* | This work |
| JDW2963 | JDW2144 *lacA::P<sub>rmB</sub>-GFPns amyE::P<sub>ilvB</sub>-mCherry* | This work |
| JDW3013 | 3610 $\Delta zpdN \Delta SP\beta \Delta PBSX \Delta comI \Delta ywaC$ | This work |
| JDW3021 | JDW2901 $\Delta yjbM \Delta ywaC \Delta relA::mIs$ | This work |
| JDW3031 | JDW2144 *guaB::pJW305 (guaB'-lacZ erm P<sub>spac</sub>-guaB)* | This work |
| JDW3517 | JDW2901 $\Delta sigE::erm$ | This work |
| JDW3519 | JDW2901 $\Delta yjbM \Delta ywaC \Delta relA::cat \Delta sigE::erm$ | This work |
| JDW3522 | YB886 $\Delta yjbM \Delta ywaC::kan \Delta relA::mIs guaB(65T\text{-}C) amyE::P_{ilvB}\text{-GFPmut2}^{A206K}$ | Bittner *et al.* 2014 and this work |
| JDW3523 | YB886 $\Delta yjbM \Delta ywaC::kan \Delta relA::mIs guaB(218A\text{-}G) amyE::P_{ilvB}\text{-GFPmut2}^{A206K}$ | Bittner *et al.* 2014 and this work |
| JDW3524 | YB886 $\Delta yjbM \Delta ywaC::kan \Delta relA::mIs guaB(361T\text{-}C) amyE::P_{ilvB}\text{-GFPmut2}^{A206K}$ | Bittner *et al.* 2014 and this work |
| JDW3525 | YB886 $\Delta yjbM \Delta ywaC::kan \Delta relA::mIs guaB(416C\text{-}T) amyE::P_{ilvB}\text{-GFPmut2}^{A206K}$ | Bittner *et al.* 2014 and this work |
| JDW3526 | YB886 $\Delta yjbM \Delta ywaC::kan \Delta relA::mIs guaB(761A\text{-}G) amyE::P_{ilvB}\text{-GFPmut2}^{A206K}$ | Bittner *et al.* 2014 and this work |
| JDW3527 | YB886 $\Delta yjbM \Delta ywaC::kan \Delta relA::mIs guaB(857C\text{-}T) amyE::P_{ilvB}\text{-GFPmut2}^{A206K}$ | Bittner *et al.* 2014 and this work |
| JDW3528 | YB886 $\Delta yjbM \Delta ywaC::kan \Delta relA::mIs guaB(943G\text{-}A) amyE::P_{ilvB}\text{-GFPmut2}^{A206K}$ | Bittner *et al.* 2014 and this work |
| JDW3529 | YB886 $\Delta yjbM \Delta ywaC::kan \Delta relA::mIs guaB(1276G\text{-}A) amyE::P_{ilvB}\text{-GFPmut2}^{A206K}$ | Bittner *et al.* 2014 and this work |
| JDW3530 | YB886 $\Delta yjbM \Delta ywaC::kan \Delta relA::mIs guaB(1330G\text{-}A) amyE::P_{ilvB}\text{-GFPmut2}^{A206K}$ | Bittner *et al.* 2014 and this work |
| JDW3531 | YB886 $\Delta yjbM \Delta ywaC::kan \Delta relA::mIs guaB(1448C\text{-}T) amyE::P_{ilvB}\text{-GFPmut2}^{A206K}$ | Bittner *et al.* 2014 and this work |
| JDW3989 | JDW2144 $\Delta guaA::kan$ | This work |
| JDW3990 | JDW2144 $\Delta purB::kan$ | This work |
| JDW3991 | JDW2144 $\Delta purL::kan$ | This work |
| JDW3992 | JDW2144 $\Delta purF::kan$ | This work |

**Table S2.** Plasmids used in this study

| Name | Genotype | Source |
| --- | --- | --- |
| pDR110 | *amyE::P<sub>spank</sub> amp spc* | David Rudner |
| pDR244 | *cre* + Ts origin | BGSC |
| pJW239 | pEX44/$\Delta yjbM$ *amp cat* | Lab stock |
| pJW299 | pEX44/I-SceI site *amp cat* | Lab stock |
| pJW300 | pJW239/$\Delta yjbM$ I-SceI site *amp cat* | Lab stock |
| pJW305 | pMUTIN4/ *P<sub>spac</sub>-guaB'-lacZ erm* | Lab stock |
| pJW306 | pJW299/$\Delta ywaC$ I-SceI site *amp cat* | Lab stock |
| pJW370 | pJW299/*yjbM* I-SceI site *amp cat* | Lab stock |
| pJW371 | pJW299/*relA*<sup>D264G</sup> I-SceI site *amp cat* | Lab stock |
| pJW562 | *yjbM*<sup>F42A</sup> I-SceI site *amp cat* | This work |
| pJW558 | pDR110/*amyE::P<sub>ilvB</sub>-gfp* | This work |
| pSS4332 | *oriU P<sub>amy</sub>-I-sceI kan* | Scott Stibitz |

**Table S3.**
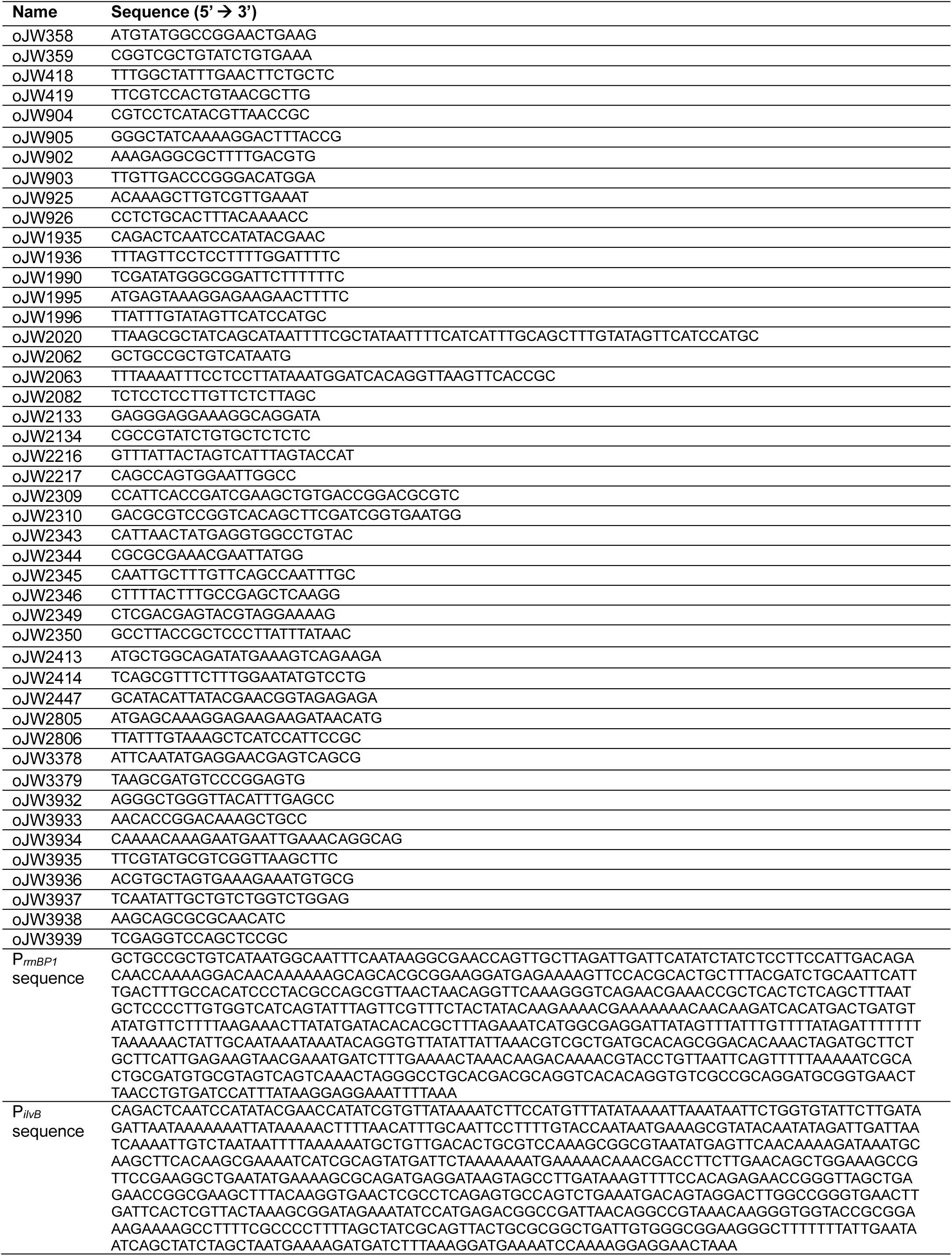
Primers and oligonucleotides used in this study

